# Nanopore-based profiling of PEGylation in nucleic acid therapeutics

**DOI:** 10.64898/2025.12.15.694309

**Authors:** Gerardo Patiño Guillén, Thieme T. Schmidt, Jeremy J. Baumberg, Jack W. Szostak, Ulrich F. Keyser, Filip Bošković

## Abstract

Nucleic acid therapeutics, including aptamers, offer effective strategies for programmable and targeted disease treatment. To improve their stability and circulation time, oligonucleotides are often conjugated to hydrophilic polymers such as polyethylene glycol (PEG). However, current bulk techniques fail to resolve PEG heterogeneity, especially in complex biological environments. Here, we use nanopore sensing to quantify PEG conjugation efficiency of the FDA-approved RNA aptamer pegaptanib. We assembled DNA nanostructures that bind pegaptanib and then we used solid-state nanopores to quantify pegaptanib PEGylation. We further assess pegaptanib PEGylation and stability in a serum background and demonstrate that nanopore sensing resolves PEG moieties of distinct molecular weights within oligonucleotide conjugates. Single-molecule profiling of polymer-RNA conjugates enables iterative improvements in oligonucleotide design and provides a direct means to assess their stability in complex biological environments, thereby advancing the development of more effective nucleic acid therapeutics.

## 1. Introduction

Nucleic acid therapeutics (NATs) offer treatment avenues for diverse diseases^1,2^ and include aptamers^3^, antisense oligonucleotides^4^, and messenger RNA therapeutics^5^, several of which have already gained FDA approval^6^. A key barrier to broader clinical adoption remains targeted and efficient delivery to specific organs or tissues^7^. Successful clinical translation requires delivery systems that enhance stability, promote strong target binding, and facilitate efficient cellular uptake^1^. To address these challenges, a variety of chemical modifications have been introduced to improve NAT stability and pharmacokinetics^7^. Particularly for shorter RNA therapeutics, PEG conjugation has proven effective, with several PEGylated NATs receiving FDA approval^8^.

PEGylation is the covalent attachment of PEG to nucleic acids at either their 3′ or 5′ ends^9^. PEG enhances solubility, reduces aggregation^10–12^, prevents renal clearance, and nuclease degradation, thereby prolonging the half-life of NATs in the bloodstream^13–16,^. PEGs with molecular weights of 20–40 kDa are commonly used in PEGylated NATs to prolong circulation time by reducing renal clearance^8,9^. PEGylation is clinically validated for RNA aptamers, including the FDA-approved pegaptanib^17^ used to treat neovascular age-related macular degeneration.

However, quantifying PEGylation and PEG’s molecular weight distribution remains challenging^18–20^. Although chromatographic methods such as HPLC are widely used for high-purity oligonucleotide analysis^21^, they often suffer from poor resolution due to polymer polydispersity or nonspecific interactions with the stationary phase^18^. Mass spectrometry is similarly limited because PEG’s flexible, neutral, and polydisperse structure produces broad, unresolved peaks and suppresses ionization^22^. These limitations are amplified in complex mixtures, preventing direct assessment of PEGylated therapeutics in biological environments. Gel electrophoresis offers a semi-quantitative tool to assess NAT PEGylation; however, its limited sensitivity and lack of sequence specificity^23^ confound quantitative analysis within complex biological backgrounds.

Although reverse transcription polymerase chain reaction (RT-PCR) is a powerful and widely used approach for RNA characterization, it has fundamental limitations for the quantitative analysis of short NATs^24,25^. Therapeutic small RNAs are often too short for direct RT-PCR because their limited sequence length constrains primer design. Assay workflows therefore commonly require adapter ligation, tailing, or other template-extension steps prior to amplification^24,25^. However, these workarounds are incompatible with chemically modified NATs where terminal modifications can prevent adapter ligation, and 2′-OH modifications can impair reverse transcription. Consequently, RT-PCR may be unsuitable for detecting chemically modified short NATs. Beyond detection, RT-PCR also provides no direct measure of PEGylation efficiency^26,27^. Addition of a radiolabel during pegaptanib synthesis or post-synthetically is an alternative but it is rarely employed due to radiation hazards^28^ and lack of labelling specificity if tritium is used^29^.

Accordingly, there is a need for a quantitative method that robustly characterises PEGylated NATs in complex biological backgrounds. Such a method should enable rapid multiplexed analysis while circumventing interference from chemical modifications, eliminating ligation-associated biases, removing the use of hazardous reagents, and supporting improved pharmacokinetic evaluation and quality control.

Here, we quantify both the degree of PEGylation and the molecular weight of PEG at the single-molecule level in NATs. Using DNA nanostructure-assisted capture^23,30^ of PEG-RNA conjugates, we characterize PEGylated NATs using nanopores^31–36^, which are compatible with PEG or PEG-containing solutions^37–41^. The specificity of nanostructure translocation enables quantitative assessment of PEGylation efficiency and multiplexed evaluation of RNA stability in complex backgrounds. Furthermore, we demonstrate that nanopores can resolve PEG molecules of distinct molecular weights within PEG mixtures, revealing polymer heterogeneity and providing a comprehensive characterization of PEGylated NATs. These capabilities establish a solid analytical framework to accelerate the development, optimization, and quality assessment of NATs.

## 2. Results and Discussion

### 2.1 DNA nanostructure-assisted capture unlocks nanopore characterization of PEGylated Nucleic Acid Therapeutics

First, we establish a strategy that combines the capture of PEGylated NATs into DNA nanostructures for their characterization by nanopore sensing. As a model therapeutic aptamer, we synthesized pegaptanib^42^ (**Figure 1a**), a 28 nt RNA aptamer of vascular endothelial growth factor (VEGF) (**Table S1**). We conjugated the 5′ end of the aptamer to PEG with a mean molecular weight of 40 kDa *via* a reaction between the 5′-amino-modifier C6 group and N-hydroxysuccinimide ester-activated PEG, and we compared it to a commercial standard (**Figure S1**).

**Figure 1.**
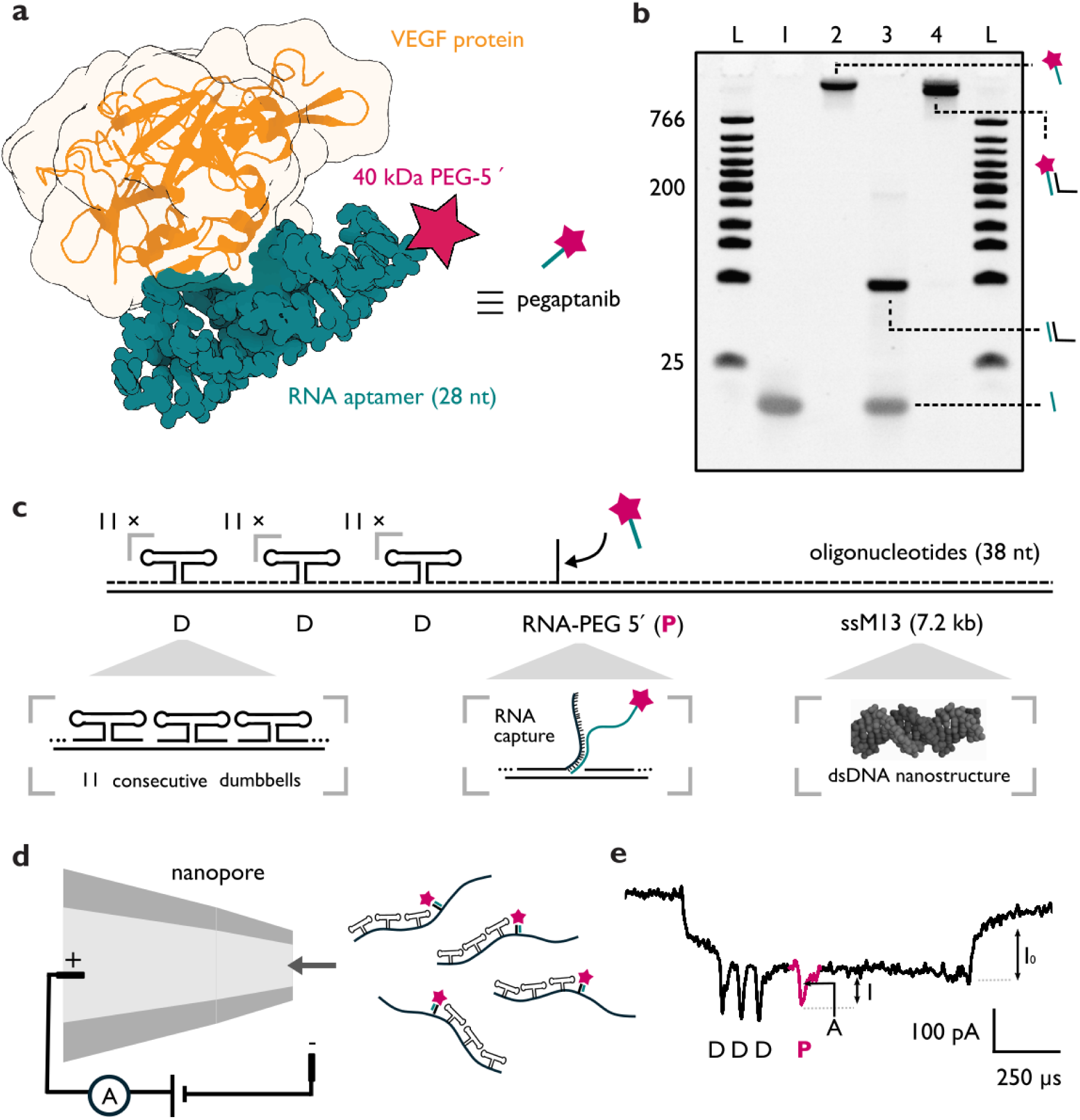
DNA nanostructure–mediated capture of pegaptanib enables PEG characterization by nanopore sensing. **a** Pegaptanib is a PEGylated (5′-40 kDa PEG) RNA aptamer that binds VEGF. **b** PAGE (10% v/v) showing hybridization of PEGylated and non-PEGylated RNA aptamer to DNA oligonucleotides: L – low-range DNA ladder, 1 – non-PEGylated RNA aptamer, 2 – PEGylated RNA aptamer, 3 – non-PEGylated RNA aptamer bound to DNA oligonucleotide (aptamer in excess), 4 – PEGylated RNA aptamer bound to DNA oligonucleotide (aptamer in excess mimicking capture conditions). **c** A 7.2 kbp dsDNA nanostructure captures pegaptanib *via* hybridization to a complementary DNA overhang. The nanostructure contains three regions with 11 dumbbells each (D), used as reference markers. **d** Nanopore-based sensing of pegaptanib-carrying nanostructure. **e** The translocation of the DNA nanostructure through the nanopore results in a drop in the ionic current. Each reference ‘D’ causes a downward current spike in the ionic current trace. Translocation of pegaptanib bound to the nanostructure produces an additional downward current spike ‘P’, enabling single-molecule characterization of the drug.

We annealed the RNA aptamer to a reverse complementary single-stranded DNA (ssDNA) oligonucleotide and performed polyacrylamide gel electrophoresis (PAGE) to demonstrate hybridization to the overhang used for nanostructure-assisted capture (**Methods**, **Figure 1b**). We hybridized both the non-PEGylated (lane 1) and PEGylated (lane 2) pegaptanib in excess to the complementary DNA oligonucleotide (lanes 3 and 4, respectively), confirming successful hybridization and RNA:DNA duplex formation (**Table S2**).

After we confirmed the hybridization of pegaptanib to the complementary oligonucleotide, we assembled a DNA nanostructure which captured pegaptanib with the same complementary oligonucleotide (**Figure 1c**). The nanostructure featured three successive references ‘D’, each composed of 11 consecutive DNA dumbbells, which provide nanopore readout selectivity. The ‘D’ regions act as barcode references. Their defined positions create a recognizable current signature that identifies the nanostructure, locates the RNA capture site, and distinguishes analyte-bearing constructs from background nucleic acids, including genomic DNA contaminants. This also enables multiplexed measurements in a single nanopore experiment^43^ (**Figure S2**; **Table S3**).

We performed nanopore ionic current measurements using the assembled DNA nanostructure. Upon the application of a voltage across a nanopore separating two chambers filled with an ionic solution, the negatively charged nanostructure, carrying pegaptanib, is driven through the pore towards the positive electrode by electrophoretic force^44^ (**Figure 1d**). As the nanostructure enters the nanopore, it displaces ions and causes a transient reduction in the ionic current *I_0_*, which is detected as the DNA nanostructure translocates through the nanopore^45,46^ (**Figure 1e**). The passage of pegaptanib through the pore induces a larger ion depletion, observed as a distinct secondary downward current spike ‘P’ with a current drop *I* and charge deficit or area *A*, enabling single-molecule quantitative detection of pegaptanib. The three references ‘D’ produce a corresponding set of three consecutive downward current spikes, providing a structural fingerprint for the nanostructure^31^ and allowing for a readout suitable for multiplexed characterization.

### 2.2 Single-molecule quantification of pegaptanib PEGylation

We quantified PEGylation of pegaptanib by simultaneous measurement of two nanostructures with distinct architectures in the same nanopore. The first nanostructure carried PEGylated pegaptanib (**Figure 2a**) and the second one bound the non-PEGylated aptamer (**Figure 2b**). To distinguish between the two constructs, the latter nanostructure was designed with two consecutive ‘D’ reference regions at one end and a third ‘D’ reference at the opposite end, generating the distinct downward current spikes during nanopore sensing and a different barcode signature (**Figure S2** and **Table S4**). When the non-PEGylated aptamer passed through the nanopore, it yielded a low prominence ‘R’ spike attributed to the 28 nt RNA:DNA hybrid between the aptamer and the nanostructure overhang. The distinct nanostructure designs allowed both constructs to translocate through the same nanopore while remaining identifiable from one another, eliminating pore-to-pore variability. This multiplexed measurement yielded two distinct current signatures within a single nanopore, the first was attributed to the nanostructure carrying the PEG-RNA, and the second to the nanostructure carrying the captured RNA, allowing direct comparison of spike observables. Both nanostructures could translocate through the pore in either the 5′→3′ or 3′→5′directions, with similar number of events in both orientations, as previously shown^47^ (**Figure 2a,b**; **Figures S3, S4**).

**Figure 2.**
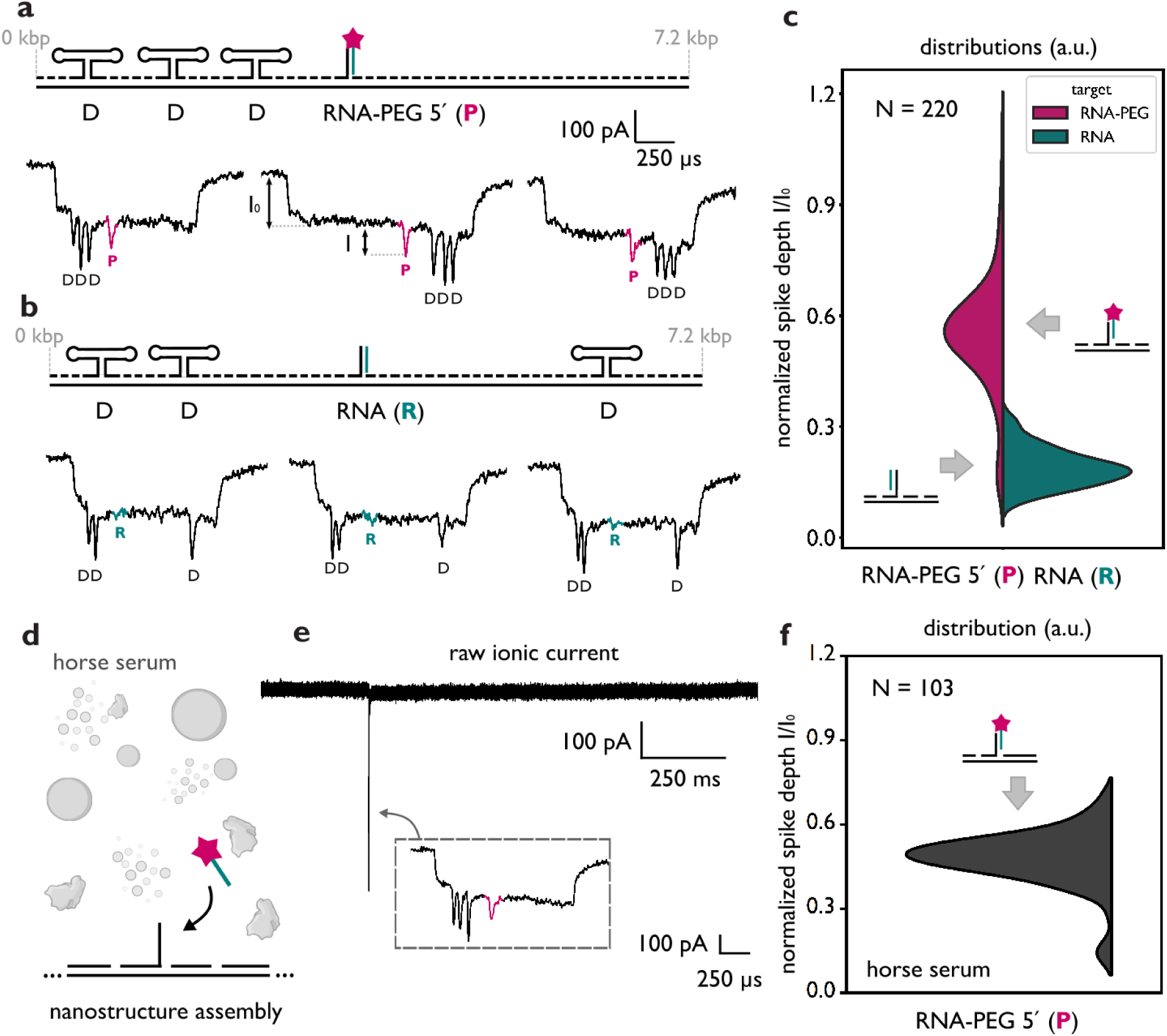
Single-molecule quantification of pegaptanib PEGylation *via* nanopore sensing. **a** We hybridized the PEGylated aptamer to a DNA nanostructure with three consecutive ‘D’ references. Translocation of this construct through a nanopore generated an initial drop in the ionic current, ascribed to the backbone of the nanostructure, followed by three consecutive downward spikes, associated with the three ‘D’ reference structures and a distinct spike ‘P’ ascribed to the 5′-40 kDa PEG RNA aptamer. **b** Non-PEGylated RNA aptamer was hybridized to a DNA nanostructure with two consecutive ‘D’ references and a terminal ‘D’ reference. Translocation of this construct through the same nanopore produced an initial drop in the ionic current associated to the backbone, two consecutive and a terminal downward current spike, associated to the three ‘D’ references and a small downward current spike ‘R’ ascribed to the RNA aptamer hybridized to the nanostructure’s overhang. **c** We compared the current spike depth of ‘P’ and ‘R’, which enabled quantitative distinction between PEGylated and non-PEGylated RNA aptamers. The mean normalized spike depth of PEGylated RNA aptamer was 0.55±0.01, while the mean depth for non-PEGylated RNA aptamer was 0.18±0.01. Errors represent the standard error of the mean (SEM). **d** We showed the detection of PEGylated pegaptanib in horse serum background. **e** Representative ionic current trace shows baseline stability and high specificity of the translocations in horse serum. **f** We observed ≈96% of PEGylation efficiency.

We measured the depth *I* of the downward current spikes ‘P’ and ‘R’, corresponding to PEGylated and non-PEGylated aptamers, respectively (**Figure 2c**), and normalized the values to *I_0_*. The spike depth depends on the nanopore’s size and geometry^48,49^, and variations in the normalized depth values are expected when measurements are performed across different nanopores. The PEGylated aptamer exhibited a mean normalized spike depth *I/I_0_* of 0.55±0.01, whereas the non-PEGylated aptamer showed a substantially lower mean value of 0.18±0.01, allowing for distinction between both aptamer species. We showed that the spike area can also be used to distinguish between the PEGylated and non-PEGylated aptamers (**Figure S5**). Notably, the normalized depth distribution of ‘P’ indicated a PEGylation level of ≈94%, which was consistent with our HPLC data (**Figure S1**). The remaining ≈6% corresponded to non-PEGylated aptamers or the translocation of an overhang which did not capture the aptamer. False-positive PEG assignments are unlikely because signals are assigned based on the position of the ‘D’ reference features and nanopore event filtering (**Figure S5**). Local knotting near the aptamer-binding site could mimic a PEG signal, but such events are expected to be rare for 7.2 kbp DNA constructs and typically shorter in duration and with higher spike amplitude^50,51^. Event-to-event variation is minimized by using internal nanostructure features to identify valid events and assigning PEG signals, with aberrant events excluded based on translocation observables and current trace profiling, as demonstrated previously^52^.

For accurate quantification, the correct design of the nanostructure overhang was essential to prevent steric hindrance by the 40 kDa PEG^9^ and thus enable efficient pegaptanib hybridization (**Figure S6**). We observed that pegaptanib binds poorly when its PEG-containing 5′ end points towards the nanostructure backbone. In contrast, when PEG is positioned away from the backbone of the nanostructure, pegaptanib shows high hybridization efficiency, as this conformation prevents steric hindrance from PEG and allows for hybridization of the RNA aptamer to the capture overhang, as depicted in **Figure 2c**.

In a different nanopore measurement, we next used PEGylated pegaptanib in horse serum as a model for monitoring the PEG-aptamer conjugates in a complex environment (**Figure 2d**). We mixed pegaptanib with serum and incubated the sample with 4 M LiCl and 3.75 M urea at 4°C for 1 hour, to disrupt intermolecular interactions of serum components and to induce the loss of protein structure^53–55^. After nanostructure assembly and nanopore characterization (**Figure 2e**), we observed consistent PEGylation efficiency (≈96%), **Figure 2f**. Raw translocations of the nanostructure and the background are presented in **Figure S7**. These results demonstrate that the nanostructure design is highly specific and can be used to assess the stability of PEG-210 based therapeutics within biological backgrounds.

### 2.3 Nanopore-based profiling of pegaptanib stability in complex backgrounds

Having demonstrated the ability to study NATs in complex biological backgrounds, we next employed nanopore sensing to monitor the stability of PEGylated aptamers in serum over time (**Figure 3**). The PEGylated aptamer and the unmodified RNA aptamer of identical sequence with a 3′ biotin, were incubated in horse serum for defined durations, followed by disruption of the serum components by treatment with 4 M LiCl and 3.75 M urea. The two species were then hybridized into structurally distinct nanostructures and measured within the same nanopore, enabling direct comparison of their stability profiles (**Figure 3a**).

**Figure 3.**
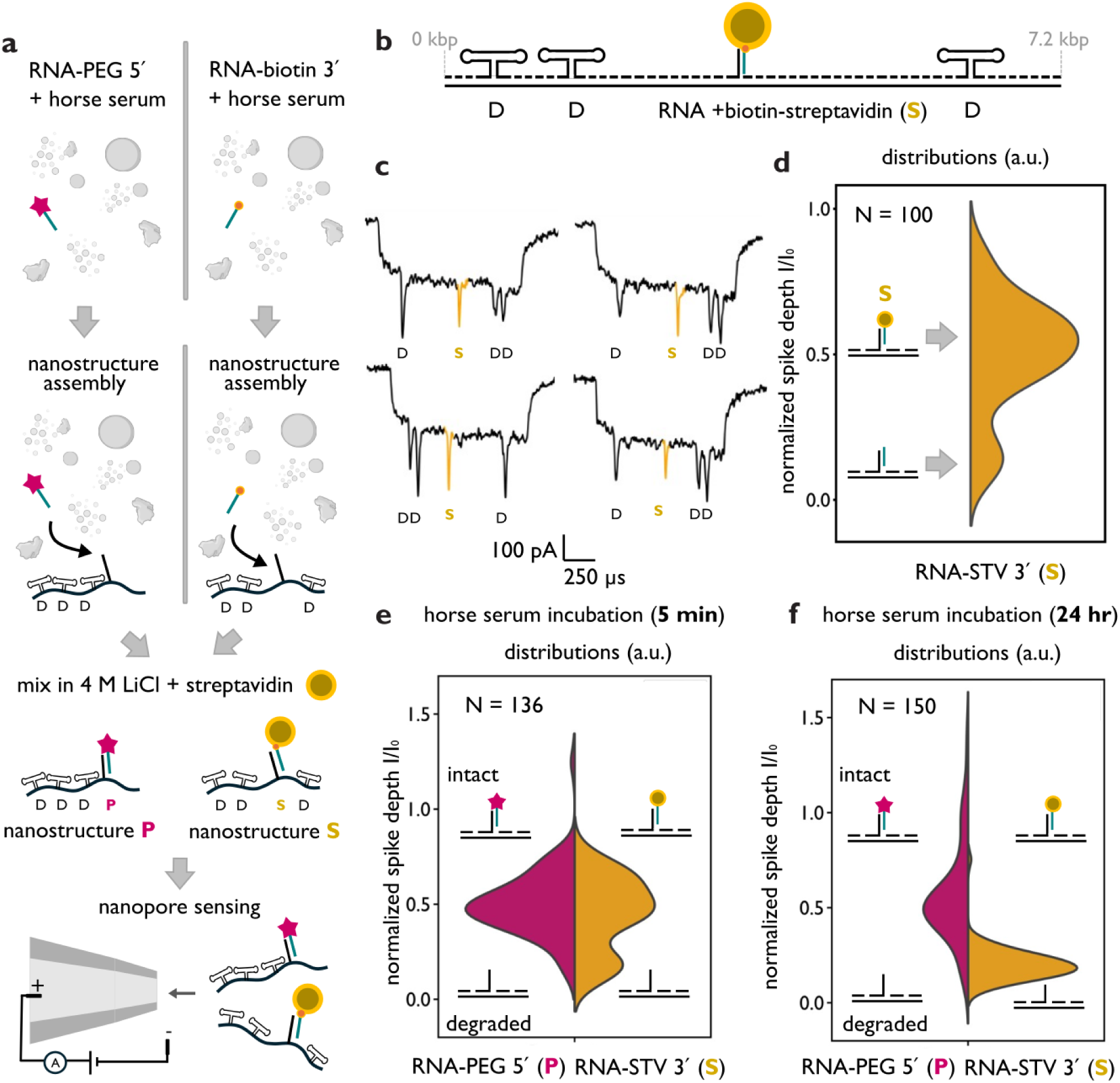
Pegaptanib degradation profiling in a complex background *via* nanopore sensing. **a** Experimental workflow in which PEGylated pegaptanib and a 3′-biotinylated RNA of identical sequence are each incubated in horse serum for defined durations, subsequently assembled into nanostructures with distinct nanopore signatures, and measured within the same nanopore for direct comparison of degradation profiles. **b** Intact biotinylated RNA is detected *via* reaction of the nanostructure with monovalent streptavidin, **c** which produces a pronounced current spike ’S’ upon translocation that serves as a spike amplification marker. **d** Streptavidin bound to intact biotinylated RNA yielded a mean normalized spike depth of 0.56±0.01. Following incubation in horse serum at 37 °C for **e** 5 min and **f** 24 h, the biotin-modified RNA aptamer displayed a progressively increasing population of translocation events in which streptavidin binding is absent, consistent with RNA degradation.

The nanostructure housing the PEGylated aptamer retained the same architecture as described for **Figure 2a**, consisting of three consecutive reference signals ’D’ followed by the PEGylated aptamer capture site ’P’. The biotinylated aptamer was incorporated into a nanostructure flanked by two consecutive ’D’ references and a terminal ’D’ reference, allowing unambiguous discrimination between the two constructs (**Figure 3b, Table S5**). Crucially, conjugation of the biotinylated aptamer nanostructure with monovalent streptavidin gave rise to a sharp current spike ’S’ during translocation which amplifies the readout of the intact biotinylated RNA aptamer (**Figure 3c**; **Figure S8**) with a mean normalized spike depth of 0.56±0.01 (**Figure 3d**).

Following incubation in horse serum at 37 °C, the PEGylated aptamer exhibited greater stability than its biotinylated counterpart. For the biotinylated aptamer, the proportion of events retaining streptavidin, which reflects intact RNA, reduced by ≈15% after 5 minutes of incubation (**Figure 3e**) and by 98% after 24 hours (**Figure 3f**), indicative of progressive RNA degradation. In contrast, the PEGylated aptamer demonstrated consistent stability across both incubation timepoints, with approximately ≈92% and ≈90% of events attributed to the intact aptamer following 5 minutes and 24 hours of incubation, respectively. The superior stability of the PEGylated aptamer is attributed to its chemical modification profile, which includes 2′-fluoro and 2′-O-methyl modifications of the ribose sugar, an inverted deoxythymidine at the 3′ end and 5′-PEGylation (**Figure S9**)^56,57^, in contrast to the 3′-biotinylated RNA aptamer lacking these modifications.

### 2.4 Determination of single-PEG polymer molecular weight

We discriminate between PEG moieties of distinct molecular weights conjugated to oligonucleotides. DNA oligonucleotides were conjugated with PEG chains of 5, 10, 20 and 40 kDa using a copper-free click reaction between 3′-dibenzocyclooctyne (DBCO) oligonucleotides and azide-functionalized PEG. Polyacrylamide gel electrophoresis (PAGE) of the PEG-conjugated oligonucleotides is shown in **Figure 4a**. As expected, increasing PEG molecular weight led to reduced migration rates (lanes 1 to 6; sequences are listed in **Table S6**).

**Figure 4.**
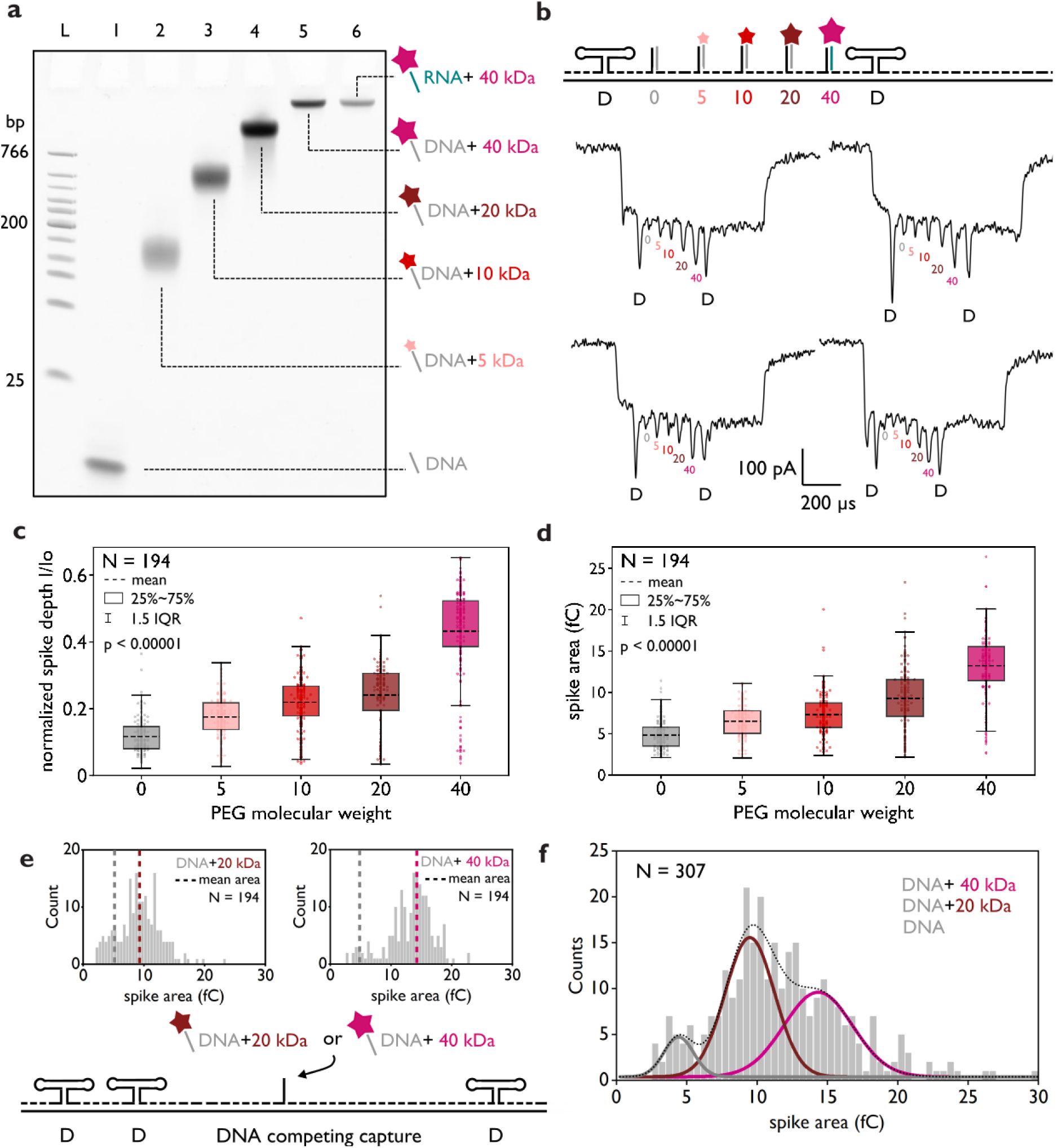
Nanopore-based determination of single-PEG polymer molar mass. **a** Native 10% PAGE shows conjugated oligonucleotides to 0 kDa (lane 1), 5 kDa (lane 2), 10 kDa (lane 3), 20 kDa (lane 4) and 40 kDa (lane 5) PEG, and the RNA aptamer conjugated to 40 kDa PEG (pegaptanib, lane 6). Lane L corresponds to a DNA ladder. **b** DNA nanostructures bearing two ‘D’ references and intervening oligonucleotides conjugated to 0, 5, 10, 20, or 40 kDa PEG produce distinct nanopore translocation events, where each PEG-containing oligonucleotide yielded a downward current spike with molecular weight-dependent **c** normalized spike depth and **d** area. **e** In the same nanopore measurement, a nanostructure with a single overhang captures an equimolar mixture of sequence-identical oligonucleotides conjugated to 20 or 40 kDa PEG. **f** Using the mean spike area measured for each pure PEG standard, we assign events in the 20/40 kDa PEG mixture to the corresponding PEG size by Gaussian fitting, thereby quantifying PEG-size heterogeneity in the sample.

The PEG-conjugated oligonucleotides were then hybridized to a unique DNA nanostructure *via* five consecutive complementary overhangs (**Figure 4b**; **Figure S2**; sequences are listed in **Table S7**). The nanostructure featured two ‘D’ references, flanking the polymer attachment sites. During nanopore sensing, an initial ionic current drop corresponded to the translocation of the nanostructure’s backbone, followed by downward spikes generated by the ‘D’ references and five additional spikes corresponding to the PEG conjugates. Both the depth and area of the polymer-associated spikes increased linearly with PEG molecular weight (**Figure S10**). Incorporation of PEG was further validated through nanopore-based detection using streptavidin binding (**Figure S11**).

PEG typically exhibits a polydispersity index (PDI) of 1.01 for molecular weights below 5 kDa, increasing to around 1.10 for PEG exceeding 50 kDa^58^. Variations in chain length affect the polymer’s flexibility and effective hydrodynamic volume, both of which influence the resulting ionic current spike-depth and area^48^. Consequently, the inherent polydispersity of PEG introduces additional variability in the measured spike observables. The normalized mean spike depths of 0, 5, 10, 20 and 40 kDa PEG were 0.12±0.01, 0.18±0.01, 0.22±0.01, 0.24±0.01 and 0.43±0.01, respectively. The error corresponds to SEM. The difference between molecular weight populations is statistically significant, showing an f-ratio value of 224.2 for a Welch’s ANOVA test and the p-value <0.00001 (**Figure 4c**). The spike areas were 4.8±0.1 fC, 6.5±0.1 fC, 7.3±0.2 fC, 9.2±0.3 fC and 13.2±0.3 fC for 0, 5, 10, 20 and 40 kDa PEG, respectively, further demonstrating we can distinguish PEG with different molecular weights (**Figure 4d**). The error corresponds to SEM, the f-ratio value was 255.1 for a Welch’s ANOVA test and the p-value <0.00001. The post-hoc Games-Howell pairwise comparisons for both observables are included in **Table S8**. Raw translocations are presented in **Figure S12**. Nanopores with diameter <5nm could be used to increase spatial resolution, allowing for discrimination of minor molecular weight variations and resolving superimposed spike prominence populations, as demonstrated previously^34,59^. The discrimination of PEG with variable molecular weight is achieved independently of the translocation direction (**Figure S13**).

In the same nanopore measurement, we included a nanostructure featuring a single capture overhang that binds an equimolar mixture of sequence-identical DNA conjugates carrying either 20 or 40 kDa PEG (**Figure 4e**). The PEG-dependent spike observables were first established from the reference nanostructure containing pure 20 and 40 kDa PEG conjugates (**Figures 4b-d**) and further compared in **Figure S14**. Among the measured observables, spike area provided the clearest separation between the 20 and 40 kDa PEG populations. We therefore used the mean spike areas of the pure 20 and 40 kDa PEG references to fit the mixed-sample distribution with a two-component Gaussian model, with each Gaussian centred on the corresponding PEG molecular weight. This analysis resolved the two PEG populations within the mixture (**Figure 4f**; raw translocation events are shown in **Figure S15**). The assignment was further validated by removing the 20 kDa PEG conjugate from the mixture, which eliminated the corresponding spike-area population (**Figure S16**).

The methodology presented demonstrates our ability to characterize oligonucleotide-polymer conjugates at the single-molecule level, enabling profiling of polymer molecular-weight distributions and identification of unexpected chain-length variants. This capability supports correlation of PEGylation profiles with therapeutic performance.

## 3. Conclusions

We have introduced a method to quantitatively study polymer-modified oligonucleotides, applying it to measure PEG levels and stability of an FDA-approved RNA aptamer. Our approach enables high-specificity multiplexed quantification of PEGylated nucleic acids in complex mixtures, such as horse serum, supporting stability assessment of PEG-based therapeutics in biological backgrounds.

The platform provides a single-molecule quantitative tool capable of analysing PEG-NAT conjugates across a broad range of molecular weights. By relying solely on nucleic acid hybridisation, it overcomes enzymatic and ligation biases, removes characterisation constraints associated with chemical modifications, offers high sensitivity and specificity, and eliminates the need for hazardous reagents. The DNA nanostructure-assisted approach provides target molecule-specific recognition^23^. In contrast, previous reports focused on the translocation of free PEGs through nanopores and did not provide the target-specific recognition of PEGylated NATs that is required for characterization in relevant complex environments^40,60^.

Beyond therapeutic monitoring, this approach is well-suited for quality control during drug manufacturing, efficacy testing, assessment of RNA stability and the rational design of drug delivery systems aimed at improving cellular uptake and pharmacokinetics. Moreover, it provides an opportunity to quantitatively assess molecular weight distribution and single-molecule behaviour of polymers with DNA nanostructure-assisted sensing, opening new directions for studying polymer–oligonucleotide conjugates.

## 4. Experimental Section

### 4.1 Pegaptanib synthesis

Oligonucleotides were synthesized using a K&A H6 oligonucleotide synthesizer, following optimized solid-phase synthesis protocols^42^. Standard and modified phosphoramidites, including 2′-F and 2′-OMe derivatives, were obtained from Glen Research or ChemGenes. To enable PEGylation, a 5′-amino-modified oligonucleotide was prepared. Subsequent conjugation to a 40-kDa N-hydroxysuccinimide (NHS)-activated PEG (purchased from MedChemExpress) was performed under standard coupling conditions and purified with 20 % (v/v) prep-scale polyacrylamide-urea gels (National Diagnostics).

### 4.2 PEG-DNA conjugates

PEG-DNA conjugates with different molecular weight were purchased from *Biomers*, their sequences are specified in **Table S6**. 5 kDa PEG-DNA was produced by request, 10 kDa: *PEG 10k (5’modification) mPEG 10kDa, mPEG 10000*, 20 kDa: *PEG 20k (5’modification) mPEG 20kDa, mPEG 20000*, 40 kDa: *PEG 40k (5’modification) mPEG 40kDa, mPEG 40000*.

### 4.3 Native polyacrylamide gel electrophoresis (PAGE)

We performed native polyacrylamide gel electrophoresis using Novex™ TBE Gels, 10% (Catalog number EC62752BOX). The constructs were run with 1 × TriTrack loading dye (Thermo Fisher Scientific, Catalog number R1161), loading 150 to 200 ng of the sample per lane. We applied a constant voltage of 80 V for 90 minutes. After electrophoresis, we stained the gels in 3 × GelRed buffer (Biotium, Catalog number 41001) and imaged using a GelDoc-It™(UVP). Gel images were processed using ImageJ (Fiji)^61^.

### 4.4 DNA nanostructure assembly

We assembled DNA nanostructures by thermal annealing. We used ssDNA M13mp18 (7,249 nt, Guild Biosciences) as scaffold. We first incubated the ssDNA with a 39 nt oligonucleotide complementary to the restriction sites of the enzymes BamHI-HF and EcoRI-HF, as reported previously^2^. We incubated ssDNA (100 nM) with the complementary oligonucleotide (2 μM) in 1 × rCutSmart Buffer (NEB). The annealing was performed at 65 °C with a linear cooling ramp to 25 °C for 40 minutes. Then, we added to the mixture 1µl of BamHI-HF (New England Biolabs (NEB), R3136S) and 1 µl of EcoRI-HF (NEB, R3101S). The reaction mixture was incubated at 37 °C for 1 hour and purified using a Monarch PCR & DNA Cleanup Kit (5 μg) (NEB, T1030S) following the manufacturer’s instructions. Then, the DNA nanostructure was assembled by mixing the cut ssDNA with the DNA oligonucleotides. The assembly reaction was performed in 10 mM MgCl_2_ (Invitrogen, AM9530G) and 1 × 10 mM Tris HCl buffer (pH 8.0). All buffers, and nuclease-free water (Ambion, Catalog number AM9937) used for assembly were previously filtered in 0.22 µm Millipore syringe filter units (MF-Merck Millipore™, Catalog number GSWP04700) and treated with UV light for 10 minutes. A reaction volume of 40 μl was implemented, which included 800 fmol of the M13 scaffold (20 nM) and a five-fold excess of DNA oligonucleotides (4 pmol, 100 nM). The target molecule for characterization (pegaptanib, or PEG-DNA conjugates) were added in a total fifteen times excess (12 pmol, 300 nM). Annealing was performed at 70 °C for 30 s, followed by cooling down over 45 min to room temperature (90 cycles of 30 s with a 0.5 °C drop per cycle). After annealing we filtered the reaction mixture twice in 100 kDa cut-off Amicon filters, to remove the excess of DNA oligonucleotides. The filtering was performed using 10 mM Tris-HCl (pH 8.0) with 0.5 mM MgCl_2_ as buffer.

### 4.5 Nanopore fabrication

We produced glass nanopores with diameters ranging from 10-15 nm^31^ using a laser-assisted capillary puller (P2000F, Sutter Instruments). The parameters of the puller were: HEAT (470-500), VEL (25), DEL (170), and PUL (200). Capillaries were purchased from Sutter Instruments and had an outer diameter of 0.5 mm and inner diameter 0.2 mm, and an inner filament. The nanopores were put into a PDMS device to facilitate measurements as described previously^52^. We performed IV curves in 4M LiCl for each nanopore produced (**Table S9**).

### 4.6 Nanopore measurements

We characterized the DNA nanostructures with nanopores. Sensing was conducted with an Axopatch 200B amplifier (Molecular Devices) in a solution containing 4 M LiCl, 1× Tris– EDTA (TE) buffer, pH 9.4. A measuring potential of 600 mV was used for translocation of the nanostructures. The amplifier held a 100 kHz internal filter, and we used an eight-pole analog low-pass Bessel filter (900CT, Frequency Devices) with a cut-off frequency of 50 kHz to filter the output signal. The data were acquired using a sampling frequency of 1 MHz in a PCI-6251 data card (National Instruments). We used custom LabVIEW codes to identify individual translocation events in the ionic current recordings. We also used custom LabVIEW codes to filter out translocation events based on duration, current drop, and charge deficit. Nonspecific translocations and drops in the ionic current were effectively removed by referencing the mean event current of the DNA nanostructure. Translocation event selection was further refined by specifying ranges of event duration and charge deficit. Unfolded translocations, where the features of the design were depicted, were selected for peak analysis. For peak characterization, we computed the spike depth and area which enabled assessment of PEGylation and PEG molecular weight.

### 4.7 DNA nanostructure assembly and characterization with horse serum background

First, we mixed 0.5 μl of horse serum (gibco, Catalog number16050-130) with 2 μl of 100 μM pegaptanib (200 pmol) for incubation periods of 0 minutes, 5 minutes and 24 hours. The mixture was diluted in 10 μl of 8 M LiCl and 7.5 μl of urea. The 20 μl mixture, holding horse serum, 10 μM pegaptanib, 4 M LiCl and 3.75 M urea was incubated for 1 hour at 4°C. The nanostructure assembly was performed in a 20 μl reaction volume containing 10 mM Tris HCl buffer (pH 8.0), 400 fmol of the M13 scaffold (20 nM) and a five-fold excess of DNA oligonucleotides (4 pmol, 100 nM). Also, 1.2 μl of serum-pegaptanib-LiCl-urea mixture was added to have a fifteen times pegaptanib excess (12 pmol, 300 nM), 240 mM LiCl and 225 mM urea. Annealing was performed at 70 °C for 30 s, followed by cooling down over 45 min to room temperature (90 cycles of 30 s with a 0.5 °C drop per cycle). After annealing we filtered the reaction mixture twice in 100 kDa cut-off Amicon filters, to remove the excess of DNA oligonucleotides. The filtering was performed using 10 mM Tris-HCl (pH 8.0) with 0.5 mM MgCl_2_ as a buffer exchange. For nanopore measurements, sample was diluted to 200 pM in 4M LiCl.

## ACKNOWLEDGMENTS

G.P.G. is supported by the Engineering and Physical Sciences Research Council (EPSRC) additional funding award for Postdoctoral Pathway Fellowships (UKRI3035), the EPSRC CDT MRes/PhD Studentship in Nanoscience and Nanotechnology (NanoDTC Cambridge EP/S022953/1), and the Trinity-Henry Barlow Scholarship. T.T.S. acknowledges UK EPSRC grant EP/X037770/1 and the European Research Council (ERC) under the Horizon2020 Research and Innovation Program PICOFORCE (883703). J.W.S. is an investigator of the Howard Hughes Medical Institute (HHMI). F.B. acknowledges funding from HHMI and the European Molecular Biology Organization through a Long-Term Fellowship (ALTF 106-2023). U.F.K. acknowledges funding from UK Research and Innovation (UKRI) under the UK government’s Horizon Europe funding guarantee EP/X023311/1, an ERC consolidator grant (DesignerPores no. 647144), an ERC-2019-PoC grant (PoreDetect no. 899538) and European Union under the Horizon2020 Program, FET-Open: DNA-FAIRYLIGHTS, Grant Agreement No. 964995. J.B. acknowledges funding from EPSRC grants EP/X037770/1, EP/Y008162/1, EP/Y036379/1 and from the ERC grant PICOFORCE 883703. We acknowledge Prof. Casey M. Platnich for insightful discussions.

## AUTHOR CONTRIBUTIONS

The manuscript was written through contributions of all authors. All authors have given approval to the final version of the manuscript. ‡These authors contributed equally.

## SUPPORTING INFORMATION

The supporting information file includes Figures S1 to S16 and Tables S1 to S9.

## TOC Graphic

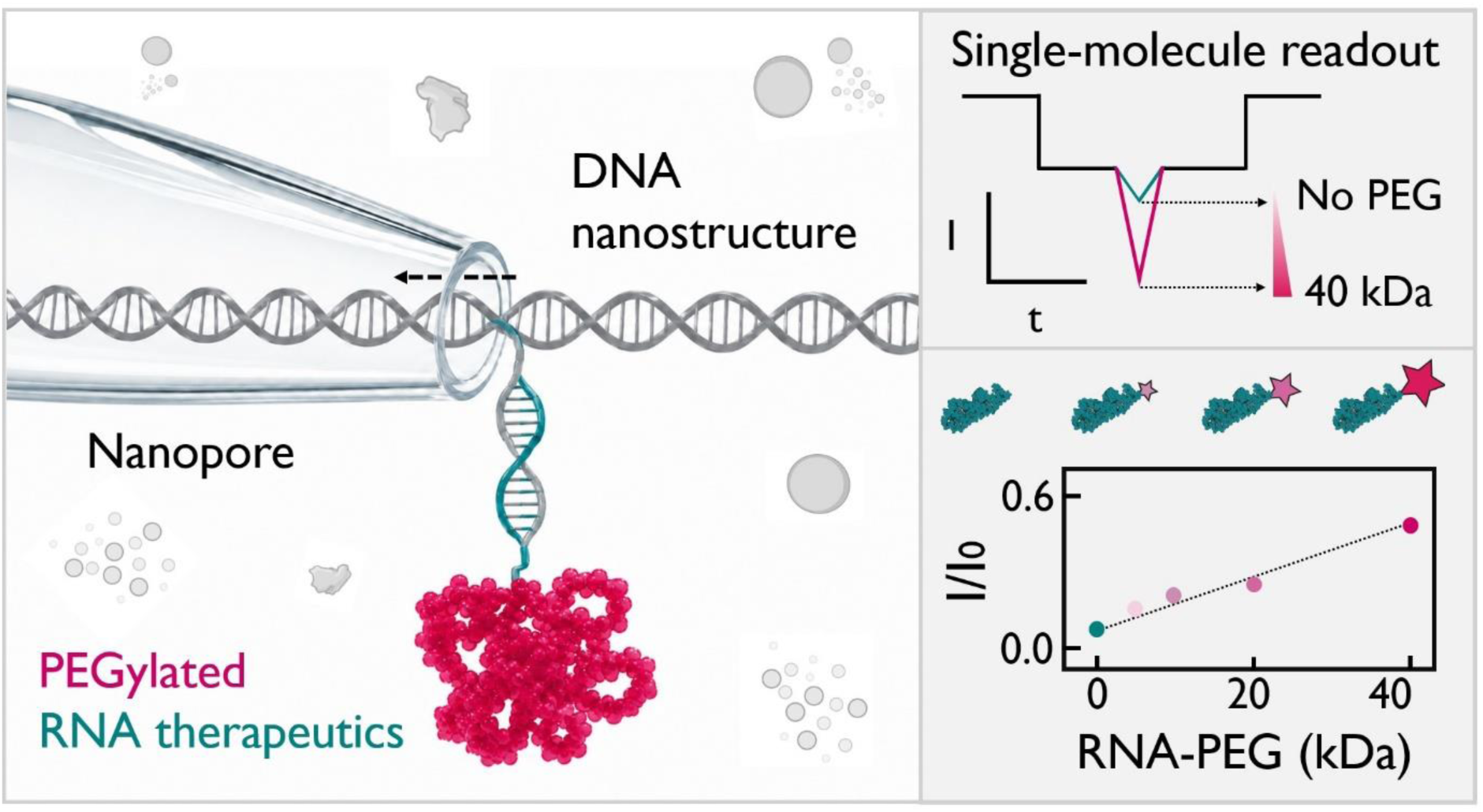

