## Supplementary Material for "Nanopore-based profiling of PEGylation in nucleic acid therapeutics"

This PDF file includes:

→ Figures S1 to S16

→ Tables S1 to S9

### Figure S1.

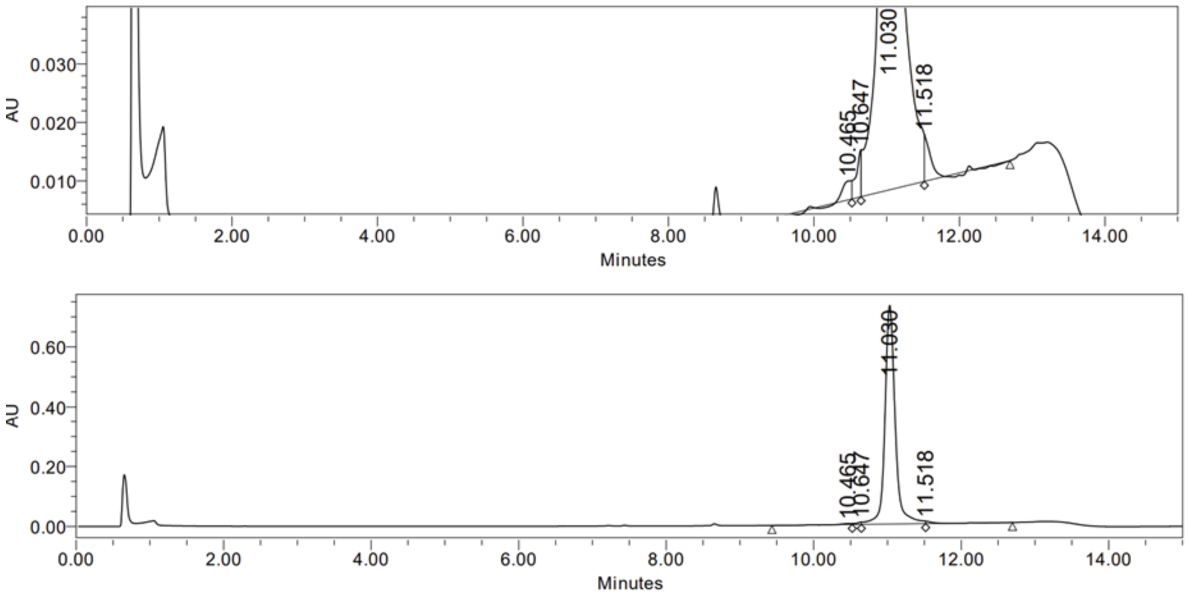


**Figure S1.** HPLC of PEGylated RNA - pegaptanib. Peak characterization indicates 98.01% of the conjugated pegaptanib, as it resolves the PEGylated RNA population from the non-PEGylated distribution.

**Figure S2.**

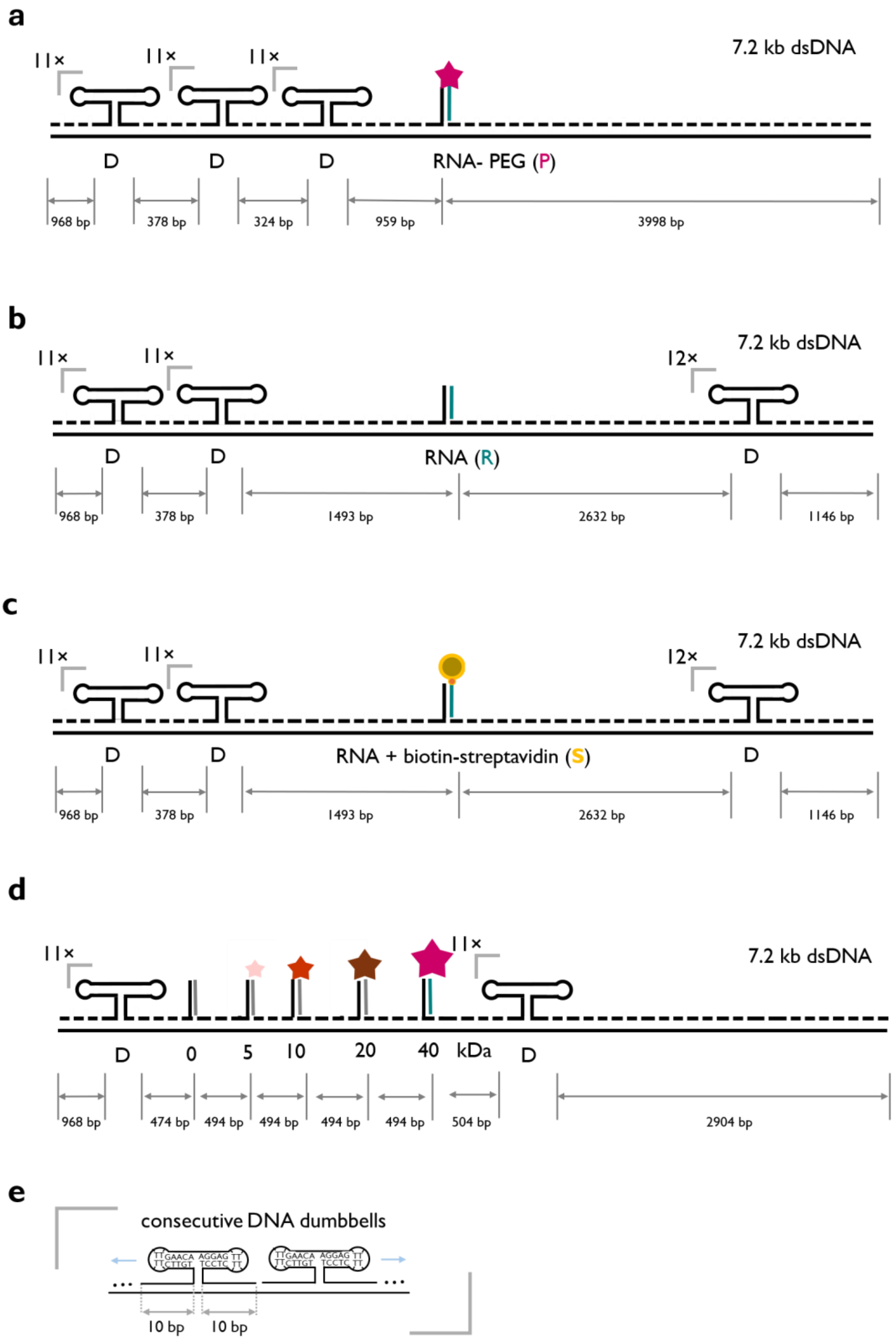


**Figure S2.** Detailed designs of DNA nanostructure holding **a** PEGylated pegaptanib ‘P’, **b** non-PEGylated pegaptanib ‘R’ and **c** biotinylated RNA ‘S’, **d** 0, 5, 10, 20 and 40 kDa PEG-oligonucleotide conjugates. **e** Detailed design of the DNA dumbbells featured in all nanostructures.

**Figure S3.**

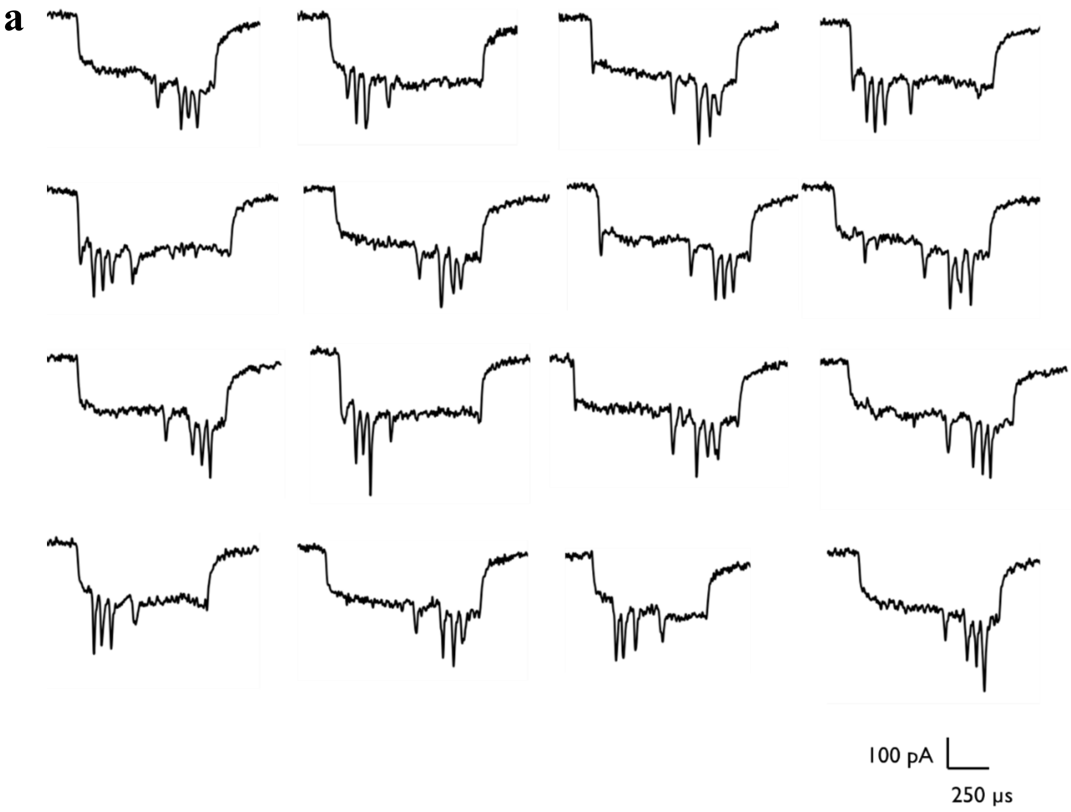


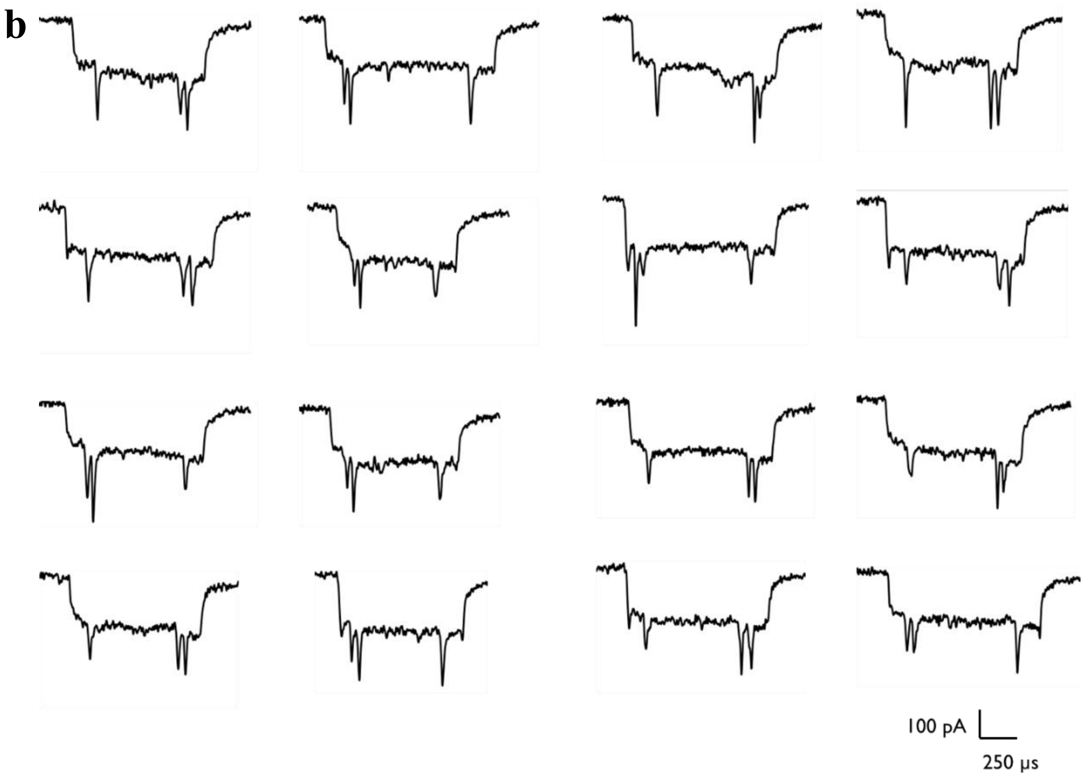


**Figure S3. a** Raw ionic current traces of unfolded DNA nanostructure holding PEGylated pegaptanib translocating through nanopore and **b** raw ionic current traces of unfolded DNA nanostructure holding RNA aptamer translocating through nanopore.

**Figure S4.**

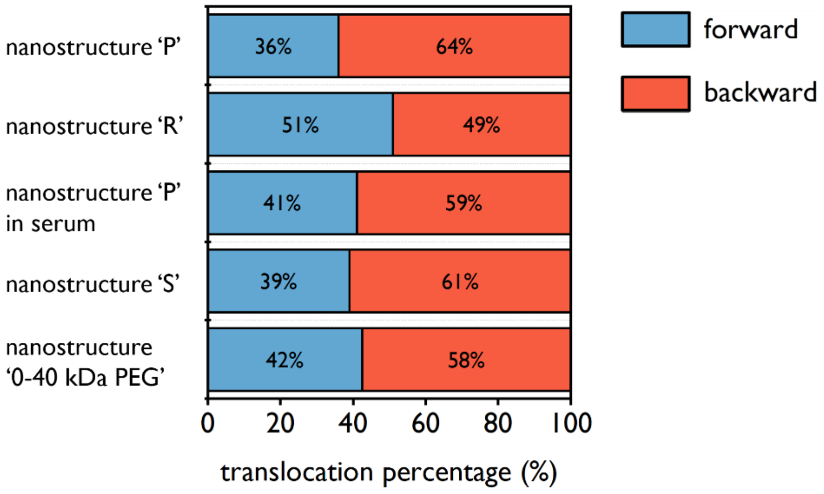


**Figure S4.** Directionality of nanostructure translocation. We show the percentage of unfolded translocation events that occur in each direction (forward 0→7.2 kbp and backward 7.2→0 kbp) for the different nanostructures sensed.

**Figure S5.**

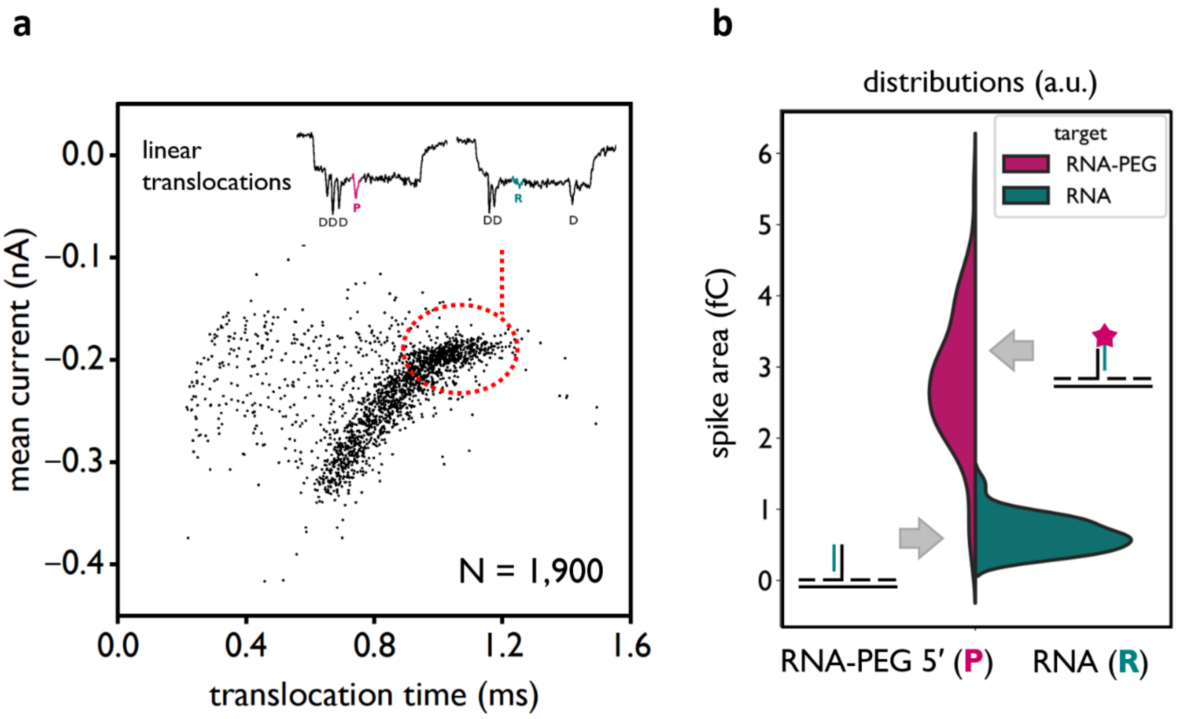


**Figure S5.** **a** Nanopore translocation observables enable the identification of linear translocation events. Unfolded nanostructures passing through the nanopore exhibit longer translocation times and smaller mean current drops. These features allow discrimination from folded translocation events, in which the nanostructure adopts a compact conformation during translocation, compromising the recognition of the nanostructure's features. **b** In linear translocation events, the current spike area is used to quantify PEGylation. We compared the current spike area of 'P' and 'R', which correspond to the spike of PEGylated and non-PEGylated pegaptanib, respectively. This enabled quantitative distinction between both aptamer species.

**Figure S6.**

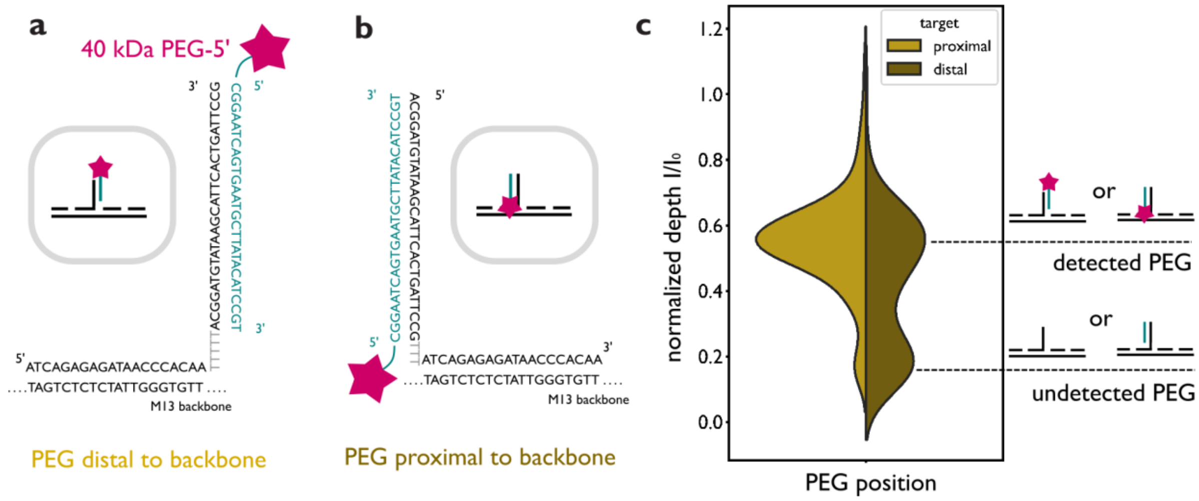


**Figure S6.** Optimization of pegaptanib capture in the DNA nanostructure. We observed that the proximity of 40 kDa PEG to the DNA nanostructure's backbone played a key role in capture efficiency. **a** Shows PEG distal to the backbone, where the PEGylated 5′ end of pegaptanib is pointing to the outside of the nanostructure and the DNA overhang has 5 'dT' as spacer. **b** Shows PEG proximal to the backbone, where the PEGylated 5′ end of pegaptanib is pointing towards the backbone and only 3 'dT' separate the overhang. **c** With the distal conformation, ≈94% of nanopore translocations showed PEGylated pegaptanib, which agrees with HPLC data. In contrast, we detected only ≈61% PEGylated pegaptanib with the proximal conformation. We attribute this difference to the steric hindrance of 40 kDa PEG to the DNA nanostructure backbone in the proximal conformation. Despite the different conformations influence capture efficiency, the normalized spike depth is not significantly affected. We utilized the distal conformation for PEGylation characterization.

### Figure S7

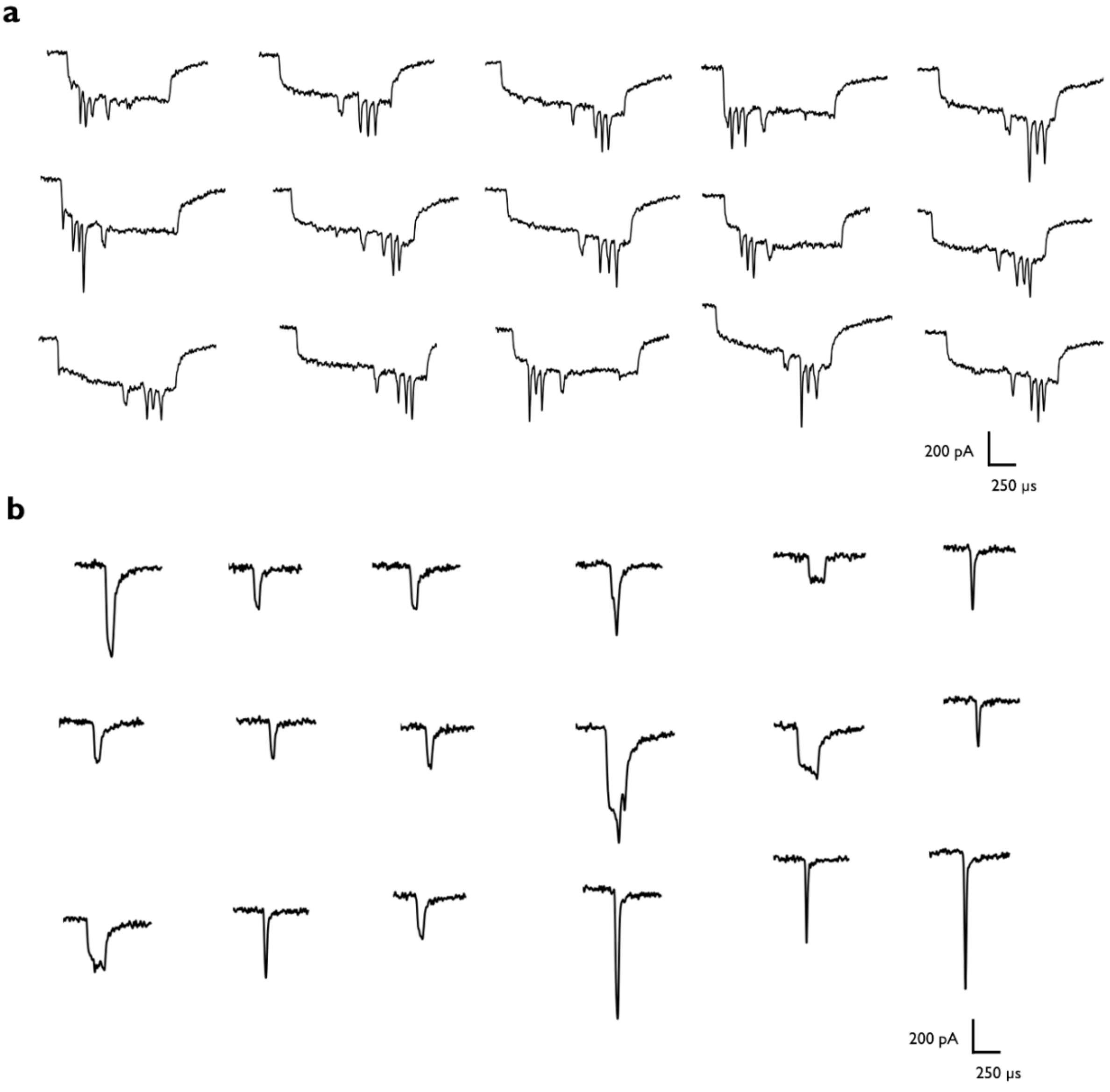


**Figure S7.** Raw nanopore ionic current traces of **a** unfolded DNA nanostructure assembled in the presence of horse serum holding PEGylated pegaptanib, and **b** non-specific translocations attributed to complex background.

**Figure S8.**

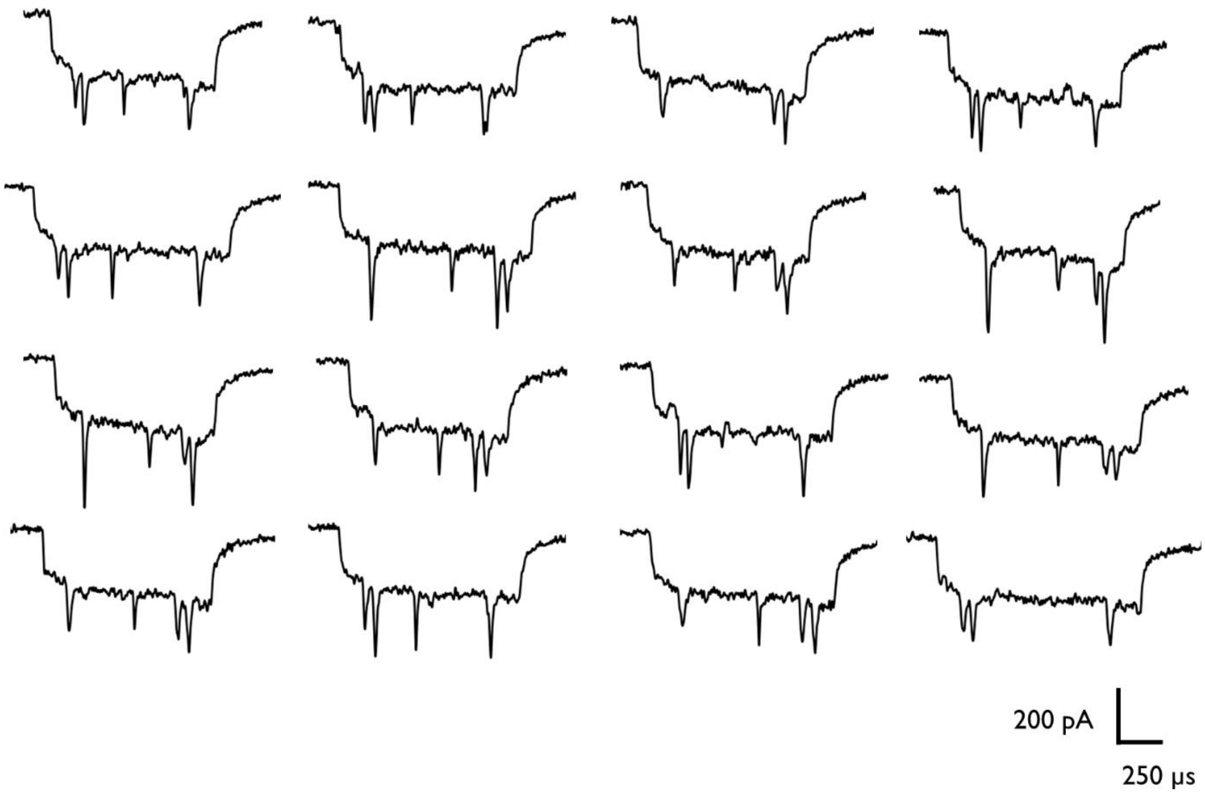


**Figure S8.** Raw nanopore ionic current traces of linear, unfolded DNA nanostructures holding RNA aptamer with a bound streptavidin-biotin complex.

**Figure S9.**

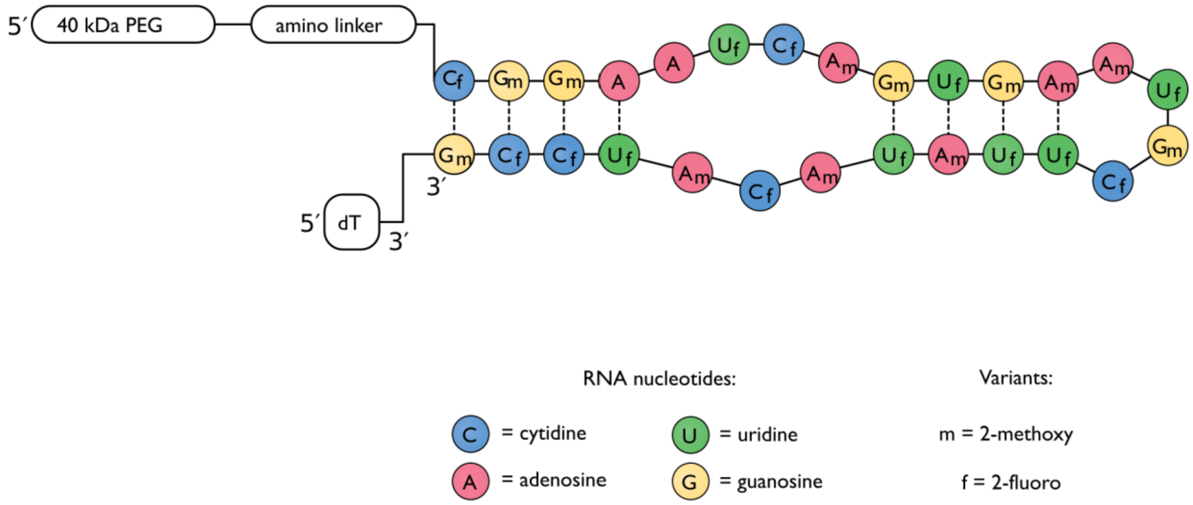


**Figure S9.** Pegaptanib sequence, indicated backbone and terminal modifications.

### Figure S10

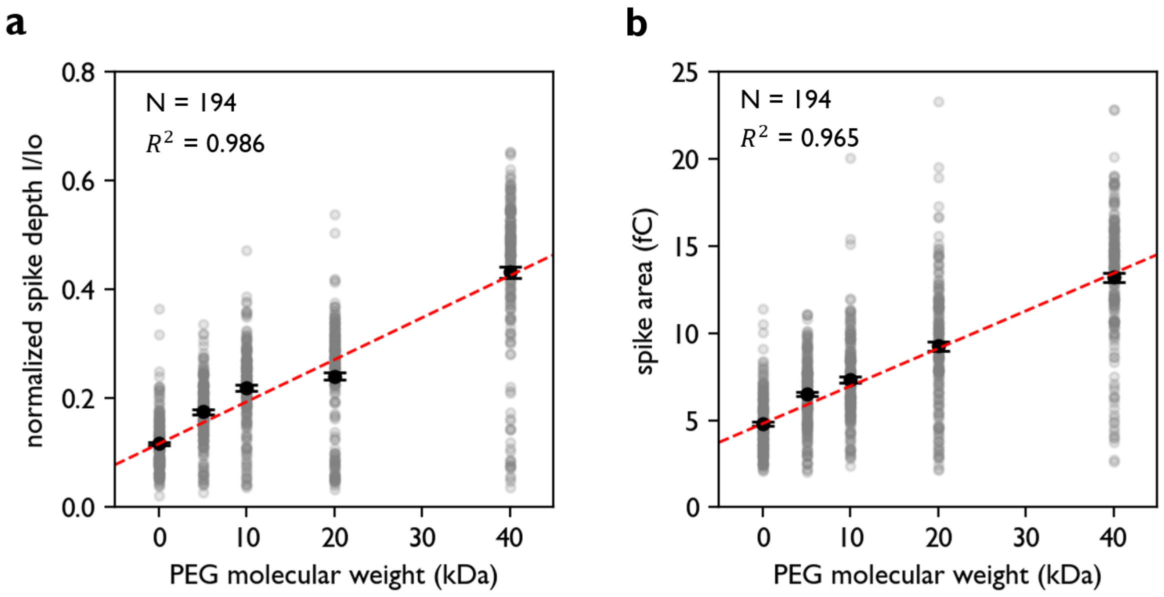


**Figure S10.** Correlation of PEG molecular weight (kDa) with **a** the normalized spike depth *I/Io* and **b** the spike area. The measurements correspond to a single nanostructure holding PEG with molecular weights of 0, 5, 10, 20 and 40 kDa.

### Figure S11.

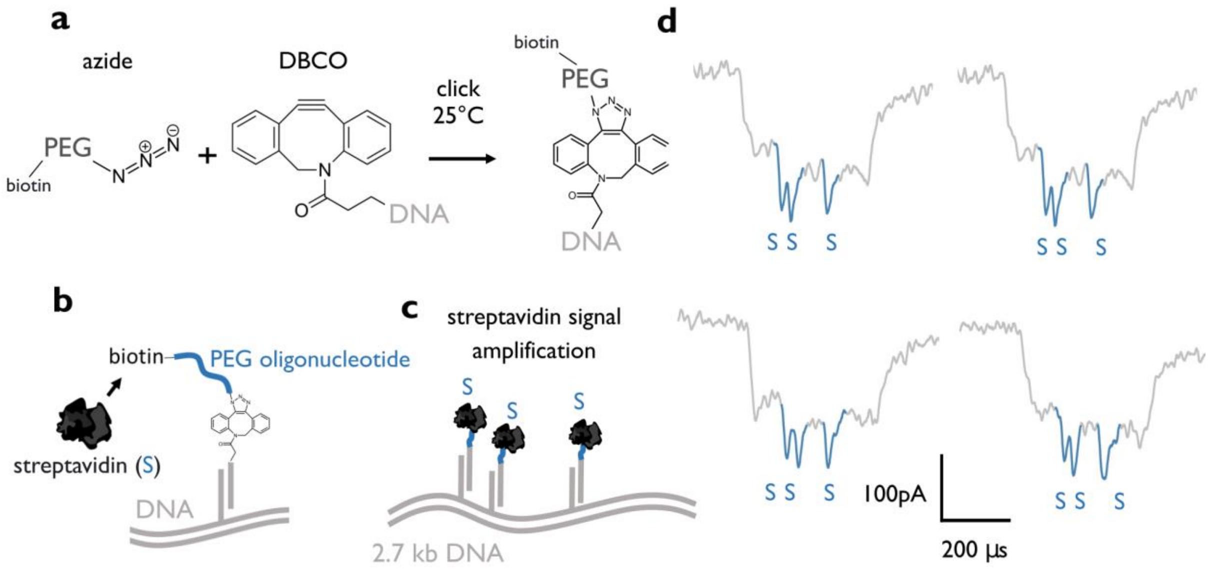


**Figure S11.** Streptavidin detection *via* biotin-PEG binding in nanopores. **a** azide-activated PEG3 oligonucleotide (Merk, 762024) and 3′-DBCO activated DNA oligonucleotide are clicked. PEG holds a biotin group in one end. **b** We capture PEG-DNA in a 2.7 kb DNA nanostructure and **c** streptavidin binds to the biotin group of PEG. **d** Streptavidin attachment produces a prominent downward current spike in nanopore sensing further validating PEG incorporation to the nanostructure.

**Figure S12.**

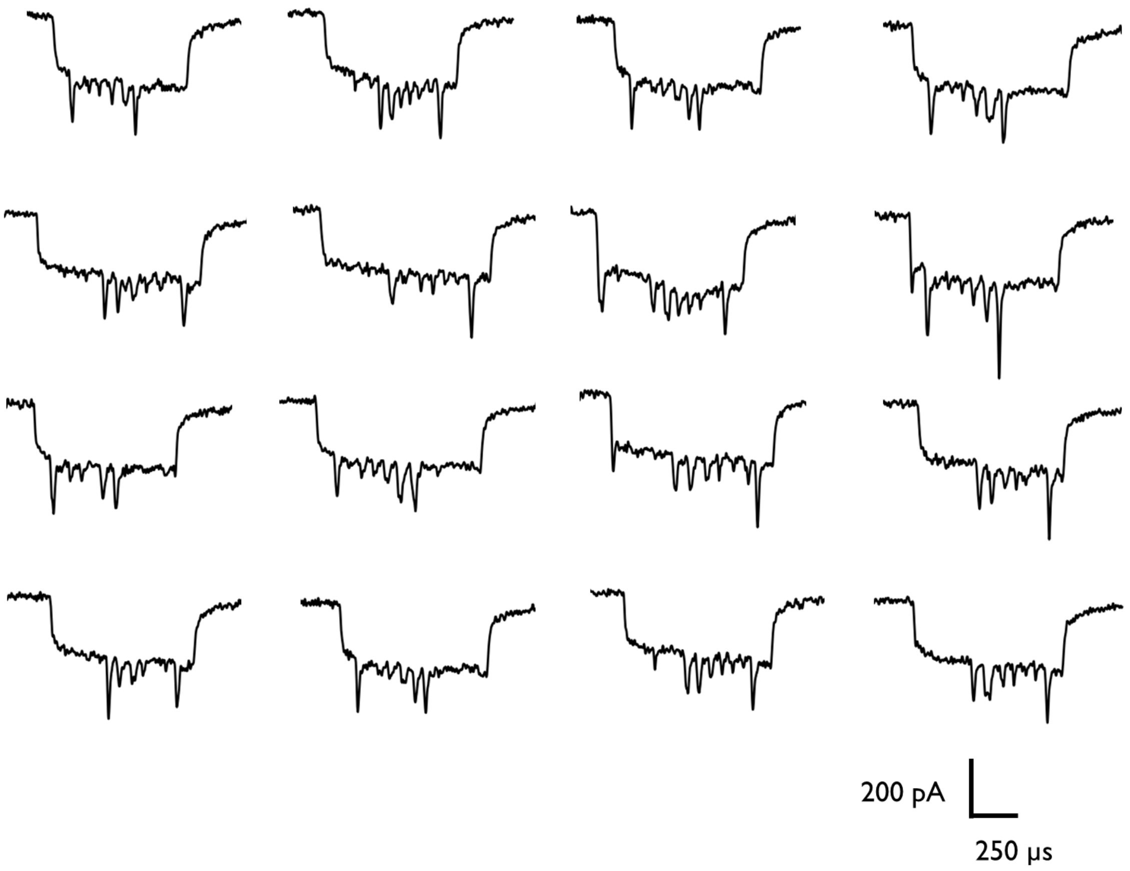


**Figure S12.** Raw ionic current traces of unfolded DNA nanostructure nanopore events holding PEGylated oligonucleotide conjugates (0,5, 10, 20 kDa) and pegaptanib (40 kDa PEG-RNA).

### Figure S13.

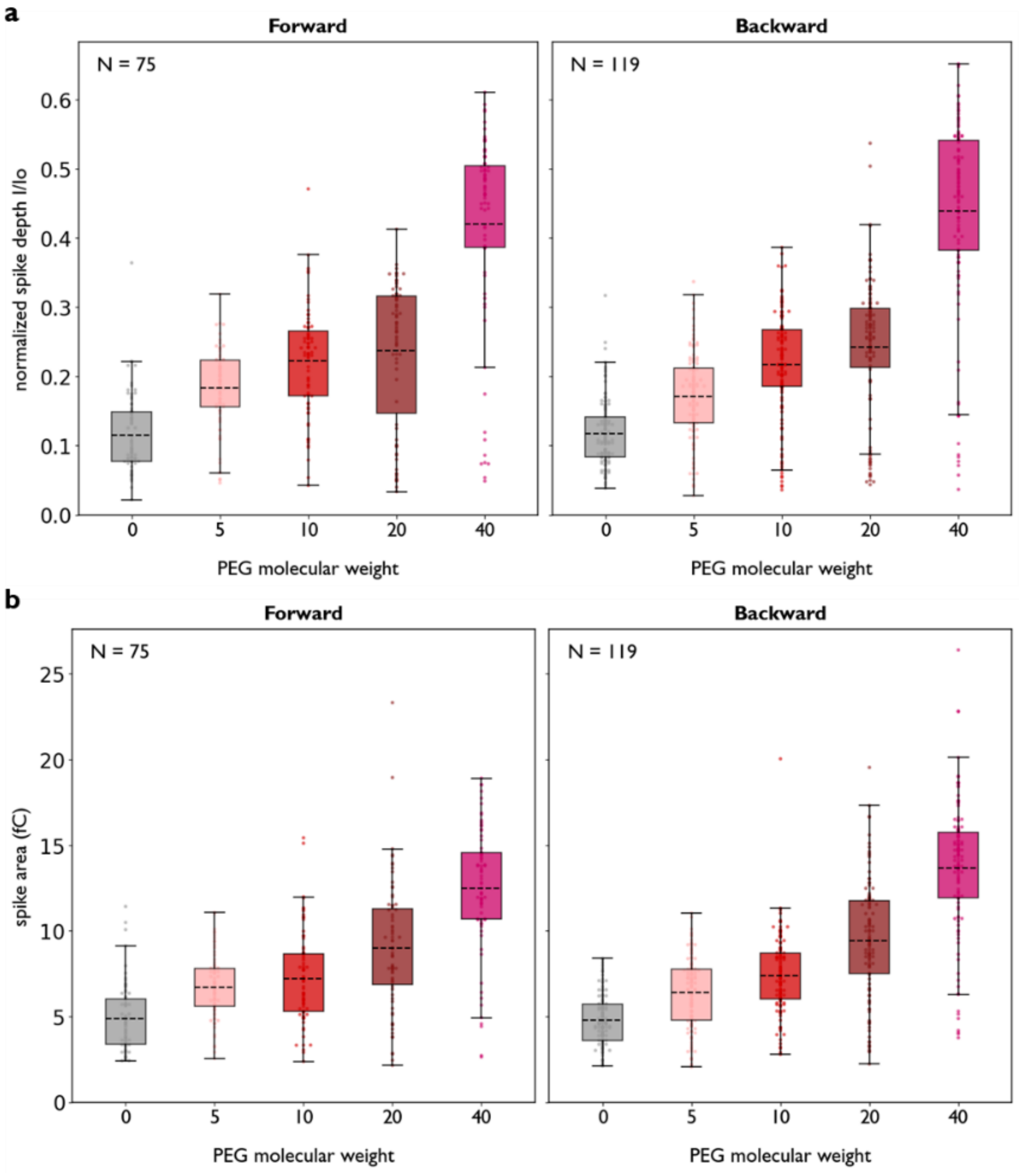


**Figure S13.** Translocation direction dependence of current spike observables. **a** Normalized spike depth for events separated by translocation direction. **b** Spike area for events separated by translocation direction. A total of 75 forward and 119 backward unfolded translocations were detected.

### Figure S14.

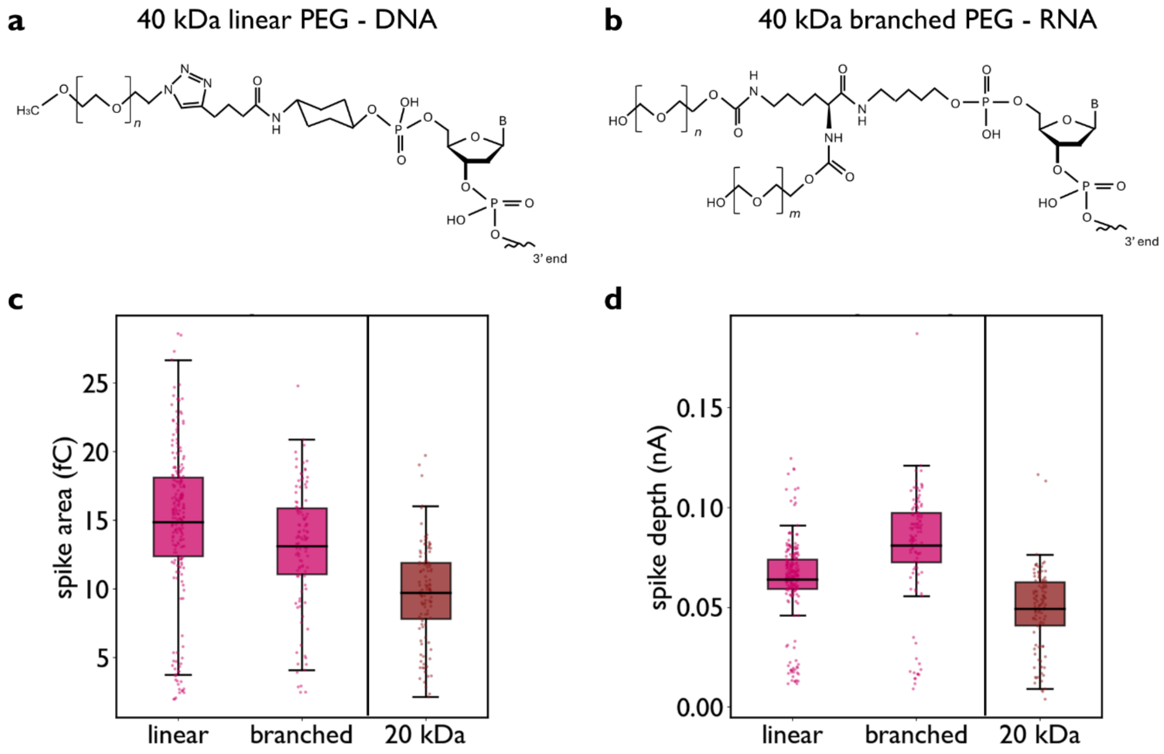


**Figure S14.** For the assessment of PEG molecular weight of in 20 and 40 kDa PEG mixtures, we use **a** 40 kDa linear PEG – DNA conjugate purchased from biomers (*mPEG 40kDa, mPEG 40000*), instead of the **b** 40 kDa branched PEG bound to the RNA aptamer (pegaptanib), to prevent hybridization biases. **c** We showed that the spike area provides better discrimination between the nanostructure holding 40 kDa linear PEG and the one with 20 kDa PEG, due to reduced overlap in the distributions.

**Figure S15.**

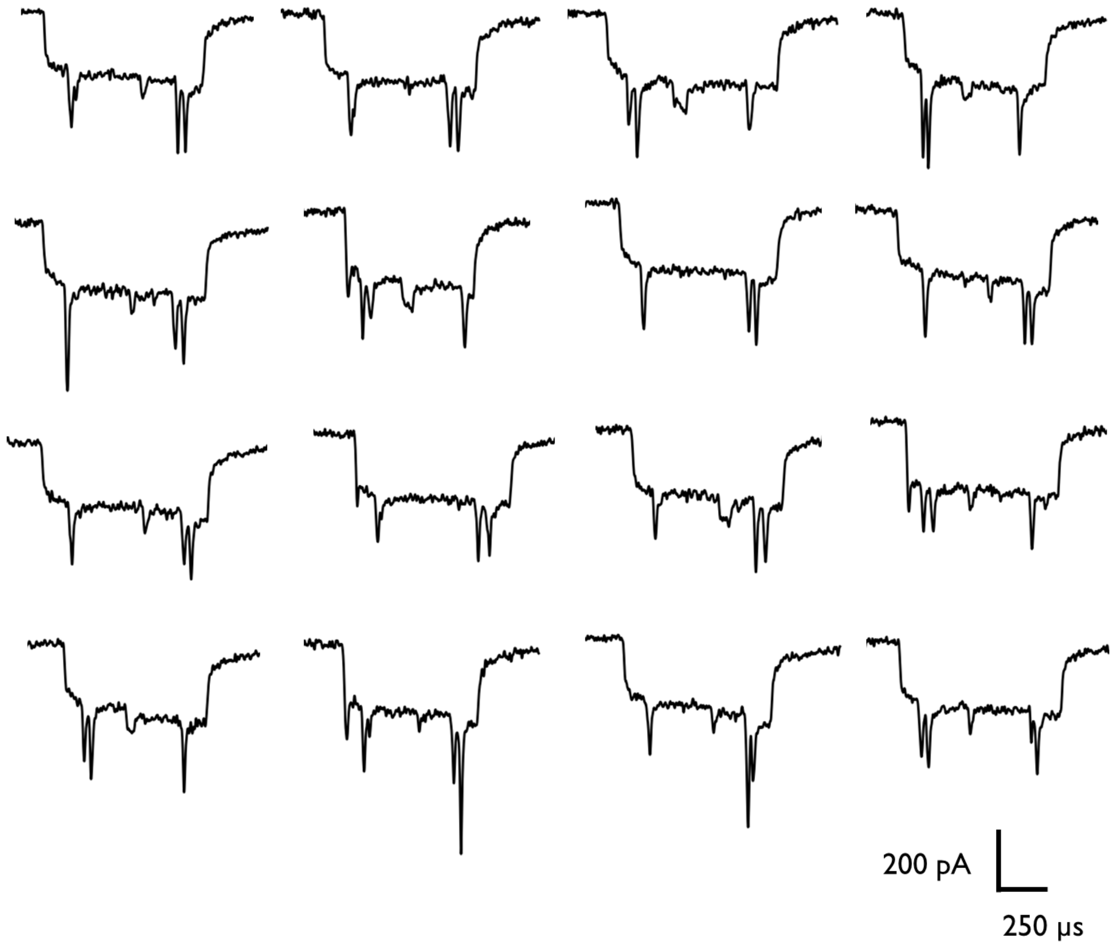


**Figure S15.** Raw nanopore ionic current traces of unfolded DNA nanostructure holding a mix of 20 and 40 kDA PEG-DNA conjugates.

### Figure S16.

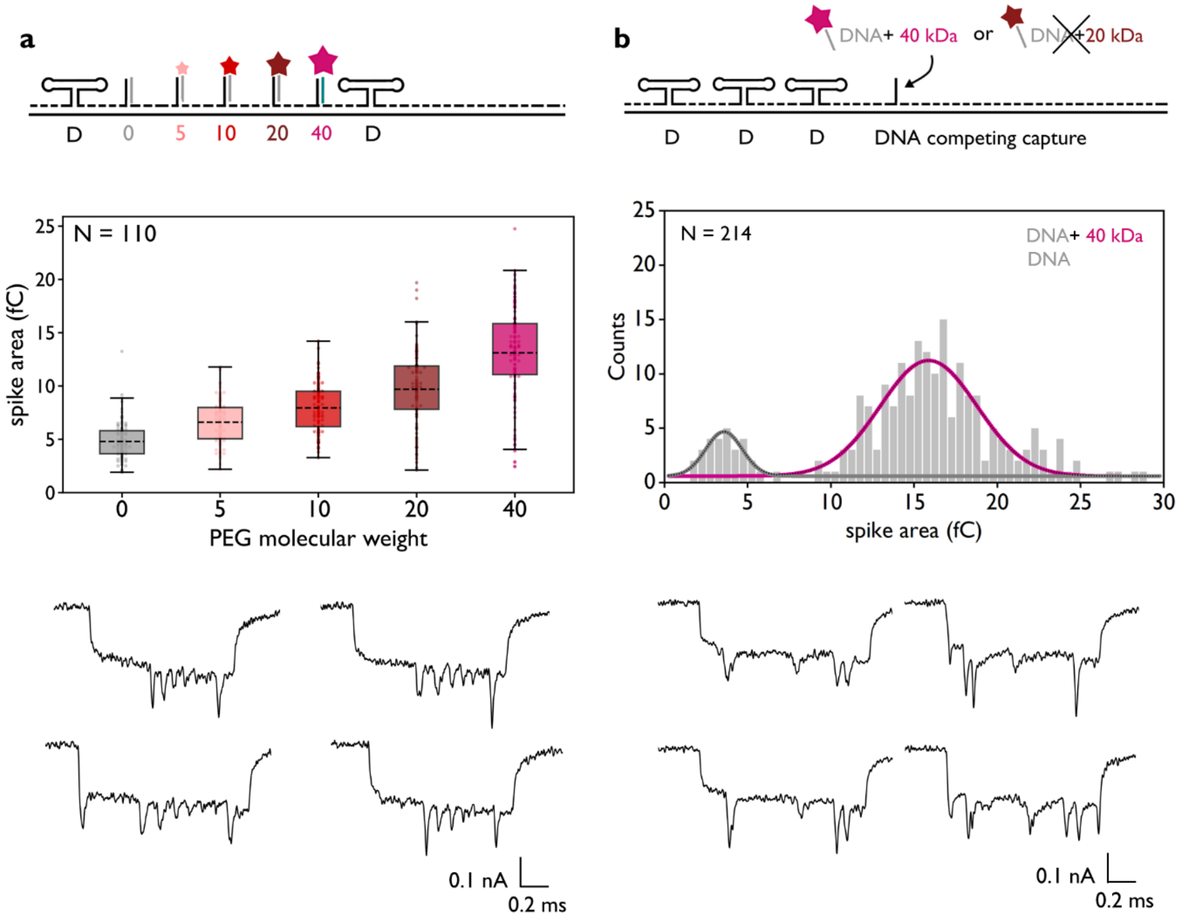


**Figure S16.** DNA nanostructure carrying 0, 5, 10, 20, and 40 kDa PEG conjugates as spike-area references for PEG molecular-weight assignment (**a**). In the same nanopore measurement, **b** we included a nanostructure which captured only the 40 kDa PEG conjugate, after removing the 20 kDa PEG conjugate from the mixture. In the spike area distribution, the population ascribed to 20 kDa PEG in Figure 4f, is no longer depicted.

### Table S1.

Sequence of PEGylated RNA aptamer, pegaptanib.

| Oligo number | Sequence (5′→ 3′) |
|---|---|
| 1 | 5′-modification (40kDa)-CGGAAUCAGUGAAUGCUUAUACAUCCGU |

### Table S2.

Sequence of 1) PEGylated RNA aptamer, 2) non-PEGylated RNA aptamer, 3) capture DNA oligonucleotide.

| Oligo number | Sequence (5′→ 3′) |
|---|---|
| 1 | 5′-modification (40kDa)-CGGAAUCAGUGAAUGCUUAUACAUCCGU |
| 2 | CGGAAUCAGUGAAUGCUUAUACAUCCGU |
| 3 | ATCAGAGAGATAACCCACAAttttttACGGATGTATAAGCATTCACTGATTCCG |

### Table S3.

Sequence of DNA oligonucleotides used for assembly of dsDNA nanostructure used for capture of PEGylated pegaptanib 'P'.

Oligonucleotide / **Dumbbell** / **Overhang**

| Oligo number | Sequence (5′→ 3′) |
|---|---|
| 1 | TTTTCGTAATCATGGTCATAGCTGTTTCCTGTGTGAAATTGTTATC |
| 2 | CGCTCACAATTCCACACAACATACGAGCCGGAAGCATA |
| 3 | AAGTGTAAAGCCTGGGGTGCCTAATGAGTGAGCTAACT |
| 4 | CACATTAATTGCGTTGCGCTCACTGCCCGCTTTCCAGT |
| 5 | CGGGAAACCTGTCGTGCCAGCTGCATTAATGAATCGGC |
| 6 | CAACGCGCGGGGAGAGGCGGTTTGCGTATTGGGCGCCA |
| 7 | GGGTGGTTTTTCTTTTCACCAGTGAGACGGGCAACAGC |
| 8 | TGATTGCCCTTCACCGCCTGGCCCTGAGAGAGTTGCAG |
| 9 | CAAGCGGTCCACGCTGGTTTGCCCCAGCAGGCGAAAAT |
| 10 | CCTGTTTGATGGTGGTTCCGAAATCGGCAAAATCCCTT |
| 11 | ATAAATCAAAAGAATAGCCCGAGATAGGGTTGAGTGTT |
| 12 | GTTCCAGTTTGGAACAAGAGTCCACTATTAAAGAACGT |
| 13 | GGACTCCAACGTCAAAGGGCGAAAAACCGTCTATCAGG |
| 14 | GCGATGGCCCACTACGTGAACCATCACCCAAATCAAGT |
| 15 | TTTTTGGGGTCGAGGTGCCGTAAAGCACTAAATCGGAA |
| 16 | CCCTAAAGGGAGCCCCCGATTTAGAGCTTGACGGGGAA |
| 17 | AGCCGGCGAACGTGGCGAGAAAGGAAGGGAAGAAAGCG |
| 18 | AAAGGAGCGGGCGCTAGGGCGCTGGCAAGTGTAGCGGT |
| 19 | CACGCTGCGCGTAACCACCACACCCGCCGCGCTTAATG |
| 20 | CGCCGCTACAGGGCGCGTACTATGGTTGCTTTGACGAG |
| 21 | CACGTATAACGTGCTTTCCTCGTTAGAATCAGAGCGGG |
| 22 | AGCTAAACAGGAGGCCGATTAAAGGGATTTTAGACAGG |
| 23 | AACGGTACGCCAGAATCCTGAGAAGTGTTTTTATAATC |
| 24 | AGTGAGGCCACCGAGTAAAAGAGTCTGTCCATCACGCA |
| 25 | AATTAACCGTTGTAGCAATACTTCTTTGATTAGTAATA |
| 26 | **ACATCACTTGTCCTCTTTTGAGGAACAAGTTTTCTTGTCCTGAGTAGA** |
| 27 | **AGAACTCAAATCCTCTTTTGAGGAACAAGTTTTCTTGTCTATCGGCCT** |
| 28 | **TGCTGGTAATTCCTCTTTTGAGGAACAAGTTTTCTTGTATCCAGAACA** |
| 29 | **ATATTACCGCTCCTCTTTTGAGGAACAAGTTTTCTTGTCAGCCATTGC** |
| 30 | **AACAGGAAAATCCTCTTTTGAGGAACAAGTTTTCTTGTACGCTCATGG** |
| 31 | **AAATACCTACTCCTCTTTTGAGGAACAAGTTTTCTTGTATTTTGACGC** |
| 32 | **TCAATCGTCTTCCTCTTTTGAGGAACAAGTTTTCTTGTGAAATGGATT** |
| 33 | **ATTTACATTGTCCTCTTTTGAGGAACAAGTTTTCTTGTGCAGATTCAC** |
| 34 | **CAGTCACACGTCCTCTTTTGAGGAACAAGTTTTCTTGTACCAGTAATA** |
| 35 | **AAAGGACATTCCTCTTTTGAGGAACAAGTTTTCTTGTTCTGGCCAAC** |

| | |
|---|---|
| 36 | **AGAGATAGAATCCTCTTTTGAGGAACAAGTTTTCTTGTCCCTTCTGAC** |
| 37 | CTGAAAGCGTAAGAATACGTGGCACAGACAATATTTTTGAATGGCT |
| 38 | ATTAGTCTTTAATGCGCGAACTGATAGCCCTAAAACAT |
| 39 | CGCCATTAAAAATACCGAACGAACCACCAGCAGAAGAT |
| 40 | AAAACAGAGGTGAGGCGGTCAGTATTAACACCGCCTGC |
| 41 | AACAGTGCCACGCTGAGAGCCAGCAGCAAATGAAAAAT |
| 42 | CTAAAGCATCACCTTGCTGAACCTCAAATATCAAACCC |
| 43 | TCAATCAATATCTGGTCAGTTGGCAAATCAACAGTTGA |
| 44 | AAGGAATTGAGGAAGGTTATCTAAAATATCTTTAGGAG |
| 45 | AATATAATCCTGATTGTTTGGATTATACTTCTGAATAATGGAAGGG |
| 46 | **CACTAACAACTCCTCTTTTGAGGAACAAGTTTTCTTGTTAATAGATTA** |
| 47 | **GAGCCGTCAATCCTCTTTTGAGGAACAAGTTTTCTTGTTAGATAATAC** |
| 48 | **ATTTGAGGATTCCTCTTTTGAGGAACAAGTTTTCTTGTTTAGAAGTAT** |
| 49 | **TAGACTTTACTCCTCTTTTGAGGAACAAGTTTTCTTGTAAACAATTCG** |
| 50 | **ACAACTCGTATCCTCTTTTGAGGAACAAGTTTTCTTGTTTAAATCCTT** |
| 51 | **TGCCCGAACGTCCTCTTTTGAGGAACAAGTTTTCTTGTTTATTAATTT** |
| 52 | **TAAAAGTTTGTCCTCTTTTGAGGAACAAGTTTTCTTGTAGTAACATTA** |
| 53 | **TCATTTTGCGTCCTCTTTTGAGGAACAAGTTTTCTTGTGAACAAAGAA** |
| 54 | **ACCACCAGAATCCTCTTTTGAGGAACAAGTTTTCTTGTGGAGCGGAAT** |
| 55 | **TATCATCATATCCTCTTTTGAGGAACAAGTTTTCTTGTTTCCTGATTA** |
| 56 | **TCAGATGATGTCCTCTTTTGAGGAACAAGTTTTCTTGTGCAATTCATC** |
| 57 | TTAGAACCTACCATATCAAAATTATTTGCACGTAAAAC |
| 58 | AGAAATAAAGAAATTGCGTAGATTTTCAGGTTTAACGT |
| 59 | CAGATGAATATACAGTAACAGTACCTTTTACATCGGGA |
| 60 | GAAACAATAACGGATTCGCCTGATTGCTTTGAATACCA |
| 61 | AGTTACAAAATCGCGCAGAGGCGAATTATTCATTTCAA |
| 62 | TTACCTGAGCAAAAGAAGATGATGAAACAAACATCAAG |
| 63 | AAAACAAAATTAATTACATTTAACAATTTCATTTGAAT |
| 64 | TACCTTTTTTAATGGAAACAGTACATAAATCAATATAT |
| 65 | **GTGAGTGAATTCCTCTTTTGAGGAACAAGTTTTCTTGTAACCTTGCTT** |
| 66 | **CTGTAAATCGTCCTCTTTTGAGGAACAAGTTTTCTTGTTCGCTATTAA** |
| 67 | **TTAATTTTCCTCCTCTTTTGAGGAACAAGTTTTCTTGTCTTAGAATCC** |
| 68 | **TTGAAAACATTCCTCTTTTGAGGAACAAGTTTTCTTGTAGCGATAGCT** |
| 69 | **TAGATTAAGATCCTCTTTTGAGGAACAAGTTTTCTTGTCGCTGAGAAG** |
| 70 | **AGTCAATAGTTCCTCTTTTGAGGAACAAGTTTTCTTGTGAATTTATCA** |
| 71 | **AAATCATAGGTCCTCTTTTGAGGAACAAGTTTTCTTGTTCTGAGAGAC** |
| 72 | **TACCTTTTTATCCTCTTTTGAGGAACAAGTTTTCTTGTACCTCCGGCT** |
| 73 | **TAGGTTGGGTTCCTCTTTTGAGGAACAAGTTTTCTTGTTATATAACTA** |
| 74 | **TATGTAAATGTCCTCTTTTGAGGAACAAGTTTTCTTGTCTGATGCAAA** |
| 75 | **TCCAATCGCATCCTCTTTTGAGGAACAAGTTTTCTTGTAGACAAAGAA** |
| 76 | CGCGAGAAAACTTTTTCAAATATATTTTAGTTAATTTCATCTTCTG |
| 77 | ACCTAAATTTAATGGTTTGAAATACCGACCGTGTGATA |
| 78 | AATAAGGCGTTAAATAAGAATAAACACCGGAATCATAA |

| | |
|---|---|
| 79 | TTACTAGAAAAAGCCTGTTTAGTATCATATGCGTTATA |
| 80 | CAAATTCTTACCAGTATAAAGCCAACGCTCAACAGTAG |
| 81 | GGCTTAATTGAGAATCGCCATATTTAACAACGCCAACA |
| 82 | TGTAATTTAGGCAGAGGCATTTTCGAGCCAGTAATAAG |
| 83 | AGAATATAAAGTACCGACAAAAGGTAAAGTAATTCTGT |
| 84 | CCAGACGACGACAATAAACAACATGTTCAGCTAATGCA |
| 85 | GAACGCGCCTGTTTATCAACAATAGATAAGTCCTGAAC |
| 86 | AAGAAAAATAATATCCCATCCTAATTTACGAGCATGTA |
| 87 | GAAACCAATCAATAATCGGCTGTCTTTCCTTATCATTC |
| 88 | CAAGAACGGGTATTAAACCAAGTACCGCACTCATCGAG |
| 89 | AACAAGCAAGCCGTTTTTATTTTCATCGTAGGAATCAT |
| 90 | TACCGCGCCCAATAGCAAGCAAATCAGATATAGAAGGC |
| 91 | TTATCCGGTATTCTAAGAACGCGAGGCGTTTTAGCGAA |
| 92 | CCTCCCGACTTGCGGGAGGTTTTGAAGCCTTAAATCAA |
| 93 | GATTAGTTGCTATTTTGCACCCAGCTACAATTTTATCC |
| 94 | TGAATCTTACCAACGCTAACGAGCGTCTTTCCAGAGCC |
| 95 | TAATTTGCCAGTTACAAAATAAACAGCCATATTATTTA |
| 96 | TCCCAATCCAAATAAGAAACGATTTTTTGTTTAACGTC |
| 97 | AAAAATGAAAATAGCAGCCTTTACAGAGAGAATAACAT |
| 98 | AAAAACAGGGAAGCGCATTAGACGGGAGAATTAACTGA |
| 99 | ACACCCTGAACAAAGTCAGAGGGTAATTGAGCGCTAAT |
| 100 | GAATTGAGTTAAGCCCAA |
| 101 | **ATCAGAGAGATAACCCACAAttttACGGATGTATAAGCATTCACTGATTCCG** |
| 102 | TAATAAGAGCAAGAAACAATGAAATAGCAATAGCTATC |
| 103 | TTACCGAAGCCCTTTTTAAGAAAAGTAAGCAGATAGCC |
| 104 | GAACAAAGTTACCAGAAGGAAACCGAGGAAACGCAATA |
| 105 | ATAACGGAATACCCAAAAGAACTGGCATGATTAAGACT |
| 106 | CCTTATTACGCAGTATGTTAGCAAACGTAGAAAATACA |
| 107 | TACATAAAGGTGGCAACATATAAAAGAAACGCAAAGAC |
| 108 | ACCACGGAATAAGTTTATTTTGTCACAATCAATAGAAA |
| 109 | ATTCATATGGTTTACCAGCGCCAAAGACAAAAGGGCGA |
| 110 | CATTCAACCGATTGAGGGAGGGAAGGTAAATATTGACG |
| 111 | GAAATTATTCATTAAAGGTGAATTATCACCGTCACCGA |
| 112 | CTTGAGCCATTTGGGAATTAGAGCCAGCAAAATCACCA |
| 113 | GTAGCACCATTACCATTAGCAAGGCCGGAAACGTCACC |
| 114 | AATGAAACCATCGATAGCAGCACCGTAATCAGTAGCGA |
| 115 | CAGAATCAAGTTTGCCTTTAGCGTCAGACTGTAGCGCG |
| 116 | TTTTCATCGGCATTTTCGGTCATAGCCCCCTTATTAGC |
| 117 | GTTTGCCATCTTTTCATAATCAAAATCACCGGAACCAG |
| 118 | AGCCACCACCGGAACCGCCTCCCTCAGAGCCGCCACCC |
| 119 | TCAGAACCGCCACCCTCAGAGCCACCACCCTCAGAGCC |
| 120 | GCCACCAGAACCACCACCAGAGCCGCCGCCAGCATTGA |
| 121 | CAGGAGGTTGAGGCAGGTCAGACGATTGGCCTTGATAT |

| | |
|---|---|
| 122 | TCACAAACAAATAAATCCTCATTAAAGCCAGAATGGAA |
| 123 | AGCGCAGTCTCTGAATTTACCGTTCCAGTAAGCGTCAT |
| 124 | ACATGGCTTTTGATGATACAGGAGTGTACTGGTAATAA |
| 125 | GTTTTAACGGGGTCAGTGCCTTGAGTAACAGTGCCCGT |
| 126 | ATAAACAGTTAATGCCCCCTGCCTATTTCGGAACCTAT |
| 127 | TATTCTGAAACATGAAAGTATTAAGAGGCTGAGACTCC |
| 128 | TCAAGAGAAGGATTAGGATTAGCGGGGTTTTGCTCAGT |
| 129 | ACCAGGCGGATAAGTGCCGTCGAGAGGGTTGATATAAG |
| 130 | TATAGCCCGGAATAGGTGTATCACCGTACTCAGGAGGT |
| 131 | TTAGTACCGCCACCCTCAGAACCGCCACCCTCAGAACC |
| 132 | GCCACCCTCAGAGCCACCACCCTCATTTTCAGGGATAG |
| 133 | CAAGCCCAATAGGAACCCATGTACCGTAACACTGAGTT |
| 134 | TCGTCACCAGTACAAACTACAACGCCTGTAGCATTCCA |
| 135 | CAGACAGCCCTCATAGTTAGCGTAACGATCTAAAGTTT |
| 136 | TGTCGTCTTTCCAGACGTTAGTAAATGAATTTTCTGTA |
| 137 | TGGGATTTTGCTAAACAACTTTCAACAGTTTCAGCGGA |
| 138 | GTGAGAATAGAAAGGAACAACTAAAGGAATTGCGAATA |
| 139 | ATAATTTTTTCACGTTGAAAATCTCCAAAAAAAAGGCT |
| 140 | CCAAAAGGAGCCTTTAATTGTATCGGTTTATCAGCTTG |
| 141 | CTTTCGAGGTGAATTTCTTAAACAGCTTGATACCGATA |
| 142 | GTTGCGCCGACAATGACAACAACCATCGCCCACGCATA |
| 143 | ACCGATATATTCGGTCGCTGAGGCTTGCAGGGAGTTAA |
| 144 | AGGCCGCTTTTGCGGGATCGTCACCCTCAGCAGCGAAA |
| 145 | GACAGCATCGGAACGAGGGTAGCAACGGCTACAGAGGC |
| 146 | TTTGAGGACTAAAGACTTTTTCATGAGGAAGTTTCCAT |
| 147 | TAAACGGGTAAAATACGTAATGCCACTACGAAGGCACC |
| 148 | AACCTAAAACGAAAGAGGCAAAAGAATACACTAAAACA |
| 149 | CTCATCTTTGACCCCCAGCGATTATACCAAGCGCGAAA |
| 150 | CAAAGTACAACGGAGATTTGTATCATCGCCTGATAAAT |
| 151 | TGTGTCGAAATCCGCGACCTGCTCCATGTTACTTAGCC |
| 152 | GGAACGAGGCGCAGACGGTCAATCATAAGGGAACCGAA |
| 153 | CTGACCAACTTTGAAAGAGGACAGATGAACGGTGTACA |
| 154 | GACCAGGCGCATAGGCTGGCTGACCTTCATCAAGAGTA |
| 155 | ATCTTGACAAGAACCGGATATTCATTACCCAAATCAAC |
| 156 | GTAACAAAGCTGCTCATTCAGTGAATAAGGCTTGCCCT |
| 157 | GACGAGAAACACCAGAACGAGTAGTAAATTGGGCTTGA |
| 158 | GATGGTTTAATTTCAACTTTAATCATTGTGAATTACCT |
| 159 | TATGCGATTTTAAGAACTGGCTCATTATACCAGTCAGG |
| 160 | ACGTTGGGAAGAAAAATCTACGTTAATAAAACGAACTA |
| 161 | ACGGAACAACATTATTACAGGTAGAAAGATTCATCAGT |
| 162 | TGAGATTTAGGAATACCACATTCAACTAATGCAGATAC |
| 163 | ATAACGCCAAAAGGAATTACGAGGCATAGTAAGAGCAA |
| 164 | CACTATCATAACCCTCGTTTACCAGACGACGATAAAAA |

| | |
|---|---|
| 165 | CCAAAATAGCGAGAGGCTTTTGCAAAAGAAGTTTTGCC |
| 166 | AGAGGGGGTAATAGTAAAATGTTTAGACTGGATAGCGT |
| 167 | CCAATACTGCGGAATCGTCATAAATATTCATTGAATCC |
| 168 | CCCTCAAATGCTTTAAACAGTTCAGAAAACGAGAATGA |
| 169 | CCATAAATCAAAAATCAGGTCTTTACCCTGACTATTAT |
| 170 | AGTCAGAAGCAAAGCGGATTGCATCAAAAAGATTAAGA |
| 171 | GGAAGCCCGAAAGACTTCAAATATCGCGTTTTAATTCG |
| 172 | AGCTTCAAAGCGAACCAGACCGGAAGCAAACTCCAACA |
| 173 | GGTCAGGATTAGAGAGTACCTTTAATTGCTCCTTTTGA |
| 174 | TAAGAGGTCATTTTTGCGGATGGCTTAGAGCTTAATTG |
| 175 | CTGAATATAATGCTGTAGCTCAACATGTTTTAAATATG |
| 176 | CAACTAAAGTACGGTGTCTGGAAGTTTCATTCCATATA |
| 177 | ACAGTTGATTCCCAATTCTGCGAACGAGTAGATTTAGT |
| 178 | TTGACCATTAGATACATTTCGCAAATGGTCAATAACCT |
| 179 | GTTTAGCTATATTTTCATTTGGGGCGCGAGCTGAAAAG |
| 180 | GTGGCATCAATTCTACTAATAGTAGTAGCATTAACATC |
| 181 | CAATAAATCATACAGGCAAGGCAAAGAATTAGCAAAAT |
| 182 | TAAGCAATAAAGCCTCAGAGCATAAAGCTAAATCGGTT |
| 183 | GTACCAAAAACATTATGACCCTGTAATACTTTTGCGGG |
| 184 | AGAAGCCTTTATTTCAACGCAAGGATAAAAATTTTTAG |
| 185 | AACCCTCATATATTTTAAATGCAATGCCTGAGTAATGT |
| 186 | GTAGGTAAAGATTCAAAAGGGTGAGAAAGGCCGGAGAC |
| 187 | AGTCAAATCACCATCAATATGATATTCAACCGTTCTAG |
| 188 | CTGATAAATTAATGCCGGAGAGGGTAGCTATTTTTGAG |
| 189 | AGATCTACAAAGGCTATCAGGTCATTGCCTGAGAGTCT |
| 190 | GGAGCAAACAAGAGAATCGATGAACGGTAATCGTAAAA |
| 191 | CTAGCATGTCAATCATATGTACCCCGGTTGATAATCAG |
| 192 | AAAAGCCCCAAAAACAGGAAGATTGTATAAGCAAATAT |
| 193 | TTAAATTGTAAACGTTAATATTTTGTTAAAATTCGCAT |
| 194 | TAAATTTTTGTTAAATCAGCTCATTTTTTAACCAATAG |
| 195 | GAACGCCATCAAAAATAATTCGCGTCTGGCCTTCCTGT |
| 196 | AGCCAGCTTTCATCAACATTAAATGTGAGCGAGTAACA |
| 197 | ACCCGTCGGATTCTCCGTGGGAACAAACGGCGGATTGA |
| 198 | CCGTAATGGGATAGGTCACGTTGGTGTAGATGGGCGCA |
| 199 | TCGTAACCGTGCATCTGCCAGTTTGAGGGGACGACGAC |
| 200 | AGTATCGGCCTCAGGAAGATCGCACTCCAGCCAGCTTT |
| 201 | CCGGCACCGCTTCTGGTGCCGGAAACCAGGCAAAGCGC |
| 202 | CATTCGCCATTCAGGCTGCGCAACTGTTGGGAAGGGCG |
| 203 | ATCGGTGCGGGCCTCTTCGCTATTACGCCAGCTGGCGA |
| 204 | AAGGGGGATGTGCTGCAAGGCGATTAAGTTGGGTAACG |
| 205 | CCAGGGTTTTCCCAGTCACGACGTTGTAAAACGACGGC |
| 206 | CAGTGCCAAGCTTGCATGCCTGCAGGTCGACTCTAGAGGATCTTTT |

### Table S4.

Sequence of DNA oligonucleotides used for assembly of dsDNA nanostructure used for capture of non-PEGylated pegaptanib ‘R’.

Oligonucleotide / **Dumbbell** / **Overhang**

| Oligo number | Sequence (5′→ 3′) |
|---|---|
| 1 | TTTTCGTAATCATGGTCATAGCTGTTTCCTGTGTGAAATTGTTATC |
| 2 | CGCTCACAATTCCACACAACATACGAGCCGGAAGCATA |
| 3 | AAGTGTAAAGCCTGGGGTGCCTAATGAGTGAGCTAACT |
| 4 | CACATTAATTGCGTTGCGCTCACTGCCCGCTTTCCAGT |
| 5 | CGGGAAACCTGTCGTGCCAGCTGCATTAATGAATCGGC |
| 6 | CAACGCGCGGGGAGAGGCGGTTTGCGTATTGGGCGCCA |
| 7 | GGGTGGTTTTTCTTTTCACCAGTGAGACGGGCAACAGC |
| 8 | TGATTGCCCTTCACCGCCTGGCCCTGAGAGAGTTGCAG |
| 9 | CAAGCGGTCCACGCTGGTTTGCCCCAGCAGGCGAAAAT |
| 10 | CCTGTTTGATGGTGGTTCCGAAATCGGCAAAATCCCTT |
| 11 | ATAAATCAAAAGAATAGCCCGAGATAGGGTTGAGTGTT |
| 12 | GTTCCAGTTTGGAACAAGAGTCCACTATTAAAGAACGT |
| 13 | GGACTCCAACGTCAAAGGGCGAAAAACCGTCTATCAGG |
| 14 | GCGATGGCCCACTACGTGAACCATCACCCAAATCAAGT |
| 15 | TTTTTGGGGTCGAGGTGCCGTAAAGCACTAAATCGGAA |
| 16 | CCCTAAAGGGAGCCCCCGATTTAGAGCTTGACGGGGAA |
| 17 | AGCCGGCGAACGTGGCGAGAAAGGAAGGGAAGAAAGCG |
| 18 | AAAGGAGCGGGCGCTAGGGCGCTGGCAAGTGTAGCGGT |
| 19 | CACGCTGCGCGTAACCACCACACCCGCCGCGCTTAATG |
| 20 | CGCCGCTACAGGGCGCGTACTATGGTTGCTTTGACGAG |
| 21 | CACGTATAACGTGCTTTCCTCGTTAGAATCAGAGCGGG |
| 22 | AGCTAAACAGGAGGCCGATTAAAGGGATTTTAGACAGG |
| 23 | AACGGTACGCCAGAATCCTGAGAAGTGTTTTTATAATC |
| 24 | AGTGAGGCCACCGAGTAAAAGAGTCTGTCCATCACGCA |
| 25 | AATTAACCGTTGTAGCAATACTTCTTTGATTAGTAATA |
| 26 | **ACATCACTTGTCCTCTTTTGAGGAACAAGTTTTCTTGTCCTGAGTAGA** |
| 27 | **AGAACTCAAATCCTCTTTTGAGGAACAAGTTTTCTTGTCTATCGGCCT** |
| 28 | **TGCTGGTAATTCCTCTTTTGAGGAACAAGTTTTCTTGTATCCAGAACA** |
| 29 | **ATATTACCGCTCCTCTTTTGAGGAACAAGTTTTCTTGTCAGCCATTGC** |
| 30 | **AACAGGAAAATCCTCTTTTGAGGAACAAGTTTTCTTGTACGCTCATGG** |
| 31 | **AAATACCTACTCCTCTTTTGAGGAACAAGTTTTCTTGTATTTTGACGC** |
| 32 | **TCAATCGTCTTCCTCTTTTGAGGAACAAGTTTTCTTGTGAAATGGATT** |

| | |
|---|---|
| 33 | **ATTTACATTGTCCTCTTTTGAGGAACAAGTTTTCTTGTGCAGATTCAC** |
| 34 | **CAGTCACACGTCCTCTTTTGAGGAACAAGTTTTCTTGTACCAGTAATA** |
| 35 | **AAAGGGACATTCCTCTTTTGAGGAACAAGTTTTCTTGTTCTGGCCAAC** |
| 36 | **AGAGATAGAATCCTCTTTTGAGGAACAAGTTTTCTTGTCCCTTCTGAC** |
| 37 | CTGAAAGCGTAAGAATACGTGGCACAGACAATATTTTTGAATGGCT |
| 38 | ATTAGTCTTTAATGCGCGAACTGATAGCCCTAAAACAT |
| 39 | CGCCATTAAAAATACCGAACGAACCACCAGCAGAAGAT |
| 40 | AAAACAGAGGTGAGGCGGTCAGTATTAACACCGCCTGC |
| 41 | AACAGTGCCACGCTGAGAGCCAGCAGCAAATGAAAAAT |
| 42 | CTAAAGCATCACCTTGCTGAACCTCAAATATCAAACCC |
| 43 | TCAATCAATATCTGGTCAGTTGGCAAATCAACAGTTGA |
| 44 | AAGGAATTGAGGAAGGTTATCTAAAATATCTTTAGGAG |
| 45 | AATATAATCCTGATTGTTTGGATTATACTTCTGAATAATGGAAGGG |
| 46 | **CACTAACAACTCCTCTTTTGAGGAACAAGTTTTCTTGTTAATAGATTA** |
| 47 | **GAGCCGTCAATCCTCTTTTGAGGAACAAGTTTTCTTGTTAGATAATAC** |
| 48 | **ATTTGAGGATTCCTCTTTTGAGGAACAAGTTTTCTTGTTTAGAAGTAT** |
| 49 | **TAGACTTTACTCCTCTTTTGAGGAACAAGTTTTCTTGTAAACAATTCG** |
| 50 | **ACAACTCGTATCCTCTTTTGAGGAACAAGTTTTCTTGTTTAAATCCTT** |
| 51 | **TGCCCGAACGTCCTCTTTTGAGGAACAAGTTTTCTTGTTTATTAATTT** |
| 52 | **TAAAAGTTTGTCCTCTTTTGAGGAACAAGTTTTCTTGTAGTAACATTA** |
| 53 | **TCATTTTGCGTCCTCTTTTGAGGAACAAGTTTTCTTGTGAACAAAGAA** |
| 54 | **ACCACCAGAATCCTCTTTTGAGGAACAAGTTTTCTTGTGGAGCGGAAT** |
| 55 | **TATCATCATATCCTCTTTTGAGGAACAAGTTTTCTTGTTTCCTGATTA** |
| 56 | **TCAGATGATGTCCTCTTTTGAGGAACAAGTTTTCTTGTGCAATTCATC** |
| 57 | TTAGAACCTACCATATCAAAATTATTTGCACGTAAAAC |
| 58 | AGAAATAAAGAAATTGCGTAGATTTTCAGGTTTAACGT |
| 59 | CAGATGAATATACAGTAACAGTACCTTTTACATCGGGA |
| 60 | GAAACAATAACGGATTCGCCTGATTGCTTTGAATACCA |
| 61 | AGTTACAAAATCGCGCAGAGGCGAATTATTCATTTCAA |
| 62 | TTACCTGAGCAAAAGAAGATGATGAAACAAACATCAAG |
| 63 | AAAACAAAATTAATTACATTTAACAATTTCATTTGAAT |
| 64 | TACCTTTTTTAATGGAAACAGTACATAAATCAATATAT |
| 65 | GTGAGTGAATAACCTTGCTTCTGTAAATCGTCGCTATT |
| 66 | AATTAATTTTCCCTTAGAATCCTTGAAAACATAGCGAT |
| 67 | AGCTTAGATTAAGACGCTGAGAAGAGTCAATAGTGAAT |
| 68 | TTATCAAAATCATAGGTCTGAGAGACTACCTTTTTAAC |
| 69 | CTCCGGCTTAGGTTGGGTTATATAACTATATGTAAATG |
| 70 | CTGATGCAAATCCAATCGCAAGACAAAGAACGCGAGAA |
| 71 | AACTTTTTCAAATATATTTTAGTTAATTTCATCTTCTG |
| 72 | ACCTAAATTTAATGGTTTGAAATACCGACCGTGTGATA |
| 73 | AATAAGGCGTTAAATAAGAATAAACACCGGAATCATAA |
| 74 | TTACTAGAAAAAGCCTGTTTAGTATCATATGCGTTATA |
| 75 | CAAATTCTTACCAGTATAAAGCCAACGCTCAACAGTAG |

| | |
|---|---|
| 76 | GGCTTAATTGAGAATCGCCATATTTAACAACGCCAACA |
| 77 | TGTAATTTAGGCAGAGGCATTTTCGAGCCAGTAATAAG |
| 78 | AGAATATAAAGTACCGACAAAAGGTAAAGTAATTCTGT |
| 79 | CCAGACGACGACAATAAACAACATGTTCAGCTAATGCA |
| 80 | GAACGCGCCTGTTTATCAACAATAGATAAGTCCTGAAC |
| 81 | AAGAAAAATAATATCCCATCCTAATTTACGAGCATGTA |
| 82 | GAAACCAATCAATAATCGGCTGTCTTTCCTTATCATTC |
| 83 | CAAGAACGGGTATTAAACCAAGTACCGCACTCATCGAG |
| 84 | AACAAGCAAGCCGTTTTTATTTTCATCGTAGGAATCAT |
| 85 | TACCGCGCCCAATAGCAAGCAAATCAGATATAGAAGGC |
| 86 | TTATCCGGTATTCTAAGAACGCGAGGCGTTTTAGCGAA |
| 87 | CCTCCCGACTTGCGGGAGGTTTTGAAGCCTTAAATCAA |
| 88 | GATTAGTTGCTATTTTGCACCCAGCTACAATTTTATCC |
| 89 | TGAATCTTACCAACGCTAACGAGCGTCTTTCCAGAGCC |
| 90 | TAATTTGCCAGTTACAAAATAAACAGCCATATTATTTA |
| 91 | TCCCAATCCAAATAAGAAACGATTTTTTGTTTAACGTC |
| 92 | AAAAATGAAAATAGCAGCCTTTACAGAGAGAATAACAT |
| 93 | AAAAACAGGGAAGCGCATTAGACGGGAGAATTAACTGA |
| 94 | ACACCCTGAACAAAGTCAGAGGGTAATTGAGCGCTAAT |
| 95 | GAATTGAGTTAAGCCCAA |
| 96 | **ATCAGAGAGATAACCCACAAttttTACGGATGTATAAGCATTCACTGATTCCG** |
| 97 | TAATAAGAGCAAGAAACAATGAAATAGCAATAGCTATC |
| 98 | TTACCGAAGCCCTTTTTAAGAAAAGTAAGCAGATAGCC |
| 99 | GAACAAAGTTACCAGAAGGAAACCGAGGAAACGCAATA |
| 100 | ATAACGGAATACCCAAAAGAACTGGCATGATTAAGACT |
| 101 | CCTTATTACGCAGTATGTTAGCAAACGTAGAAAATACA |
| 102 | TACATAAAGGTGGCAACATATAAAAGAAACGCAAAGAC |
| 103 | ACCACGGAATAAGTTTATTTTGTCACAATCAATAGAAA |
| 104 | ATTCATATGGTTTACCAGCGCCAAAGACAAAAGGGCGA |
| 105 | CATTCAACCGATTGAGGGAGGGAAGGTAAATATTGACG |
| 106 | GAAATTATTCATTAAAGGTGAATTATCACCGTCACCGA |
| 107 | CTTGAGCCATTTGGGAATTAGAGCCAGCAAAATCACCA |
| 108 | GTAGCACCATTACCATTAGCAAGGCCGGAAACGTCACC |
| 109 | AATGAAACCATCGATAGCAGCACCGTAATCAGTAGCGA |
| 110 | CAGAATCAAGTTTGCCTTTAGCGTCAGACTGTAGCGCG |
| 111 | TTTTCATCGGCATTTTCGGTCATAGCCCCCTTATTAGC |
| 112 | GTTTGCCATCTTTTCATAATCAAAATCACCGGAACCAG |
| 113 | AGCCACCACCGGAACCGCCTCCCTCAGAGCCGCCACCC |
| 114 | TCAGAACCGCCACCCTCAGAGCCACCACCCTCAGAGCC |
| 115 | GCCACCAGAACCACCACCAGAGCCGCCGCCAGCATTGA |
| 116 | CAGGAGGTTGAGGCAGGTCAGACGATTGGCCTTGATAT |
| 117 | TCACAAACAAATAAATCCTCATTAAAGCCAGAATGGAA |
| 118 | AGCGCAGTCTCTGAATTTACCGTTCCAGTAAGCGTCAT |

| | |
|---|---|
| 119 | ACATGGCTTTTGATGATACAGGAGTGTACTGGTAATAA |
| 120 | GTTTTAACGGGGTCAGTGCCTTGAGTAACAGTGCCCGT |
| 121 | ATAAACAGTTAATGCCCCCTGCCTATTTCGGAACCTAT |
| 122 | TATTCTGAAACATGAAAGTATTAAGAGGCTGAGACTCC |
| 123 | TCAAGAGAAGGATTAGGATTAGCGGGGTTTTGCTCAGT |
| 124 | ACCAGGCGGATAAGTGCCGTCGAGAGGGTTGATATAAG |
| 125 | TATAGCCCGGAATAGGTGTATCACCGTACTCAGGAGGT |
| 126 | TTAGTACCGCCACCCTCAGAACCGCCACCCTCAGAACC |
| 127 | GCCACCCTCAGAGCCACCACCCTCATTTTCAGGGATAG |
| 128 | CAAGCCCAATAGGAACCCATGTACCGTAACACTGAGTT |
| 129 | TCGTCACCAGTACAAACTACAACGCCTGTAGCATTCCA |
| 130 | CAGACAGCCCTCATAGTTAGCGTAACGATCTAAAGTTT |
| 131 | TGTCGTCTTTCCAGACGTTAGTAAATGAATTTTCTGTA |
| 132 | TGGGATTTTGCTAAACAACTTTCAACAGTTTCAGCGGA |
| 133 | GTGAGAATAGAAAGGAACAACTAAAGGAATTGCGAATA |
| 134 | ATAATTTTTTCACGTTGAAAATCTCCAAAAAAAAAGGCT |
| 135 | CCAAAAGGAGCCTTTAATTGTATCGGTTTATCAGCTTG |
| 136 | CTTTCGAGGTGAATTTCTTAAACAGCTTGATACCGATA |
| 137 | GTTGCGCCGACAATGACAACAACCATCGCCCACGCATA |
| 138 | ACCGATATATTCGGTCGCTGAGGCTTGCAGGGAGTTAA |
| 139 | AGGCCGCTTTTGCGGGATCGTCACCCTCAGCAGCGAAA |
| 140 | GACAGCATCGGAACGAGGGTAGCAACGGCTACAGAGGC |
| 141 | TTTGAGGACTAAAGACTTTTTCATGAGGAAGTTTCCAT |
| 142 | TAAACGGGTAAAATACGTAATGCCACTACGAAGGCACC |
| 143 | AACCTAAAACGAAAGAGGCAAAAGAATACACTAAAACA |
| 144 | CTCATCTTTGACCCCCAGCGATTATACCAAGCGCGAAA |
| 145 | CAAAGTACAACGGAGATTTGTATCATCGCCTGATAAAT |
| 146 | TGTGTCGAAATCCGCGACCTGCTCCATGTTACTTAGCC |
| 147 | GGAACGAGGCGCAGACGGTCAATCATAAGGGAACCGAA |
| 148 | CTGACCAACTTTGAAAGAGGACAGATGAACGGTGTACA |
| 149 | GACCAGGCGCATAGGCTGGCTGACCTTCATCAAGAGTA |
| 150 | ATCTTGACAAGAACCGGATATTCATTACCCAAATCAAC |
| 151 | GTAACAAAGCTGCTCATTCAGTGAATAAGGCTTGCCCT |
| 152 | GACGAGAAACACCAGAACGAGTAGTAAATTGGGCTTGA |
| 153 | GATGGTTTAATTTCAACTTTAATCATTGTGAATTACCT |
| 154 | TATGCGATTTTAAGAACTGGCTCATTATACCAGTCAGG |
| 155 | ACGTTGGGAAGAAAAATCTACGTTAATAAAACGAACTA |
| 156 | ACGGAACAACATTATTACAGGTAGAAAGATTCATCAGT |
| 157 | TGAGATTTAGGAATACCACATTCAACTAATGCAGATAC |
| 158 | ATAACGCCAAAAGGAATTACGAGGCATAGTAAGAGCAA |
| 159 | CACTATCATAACCCTCGTTTACCAGACGACGATAAAAA |
| 160 | CCAAAATAGCGAGAGGCTTTTGCAAAAGAAGTTTTGCC |
| 161 | AGAGGGGGTAATAGTAAAATGTTTAGACTGGATAGCGT |

| | |
|---|---|
| 162 | CCAATACTGCGGAATCGTCATAAATATTCATTGAATCC |
| 163 | CCCTCAAATGCTTTAAACAGTTCAGAAAACGAGAATGA |
| 164 | CCATAAATCAAAAATCAGGTCTTTACCCTGACTATTAT |
| 165 | AGTCAGAAGCAAAGCGGATTGCATCAAAAAGATTAAGA |
| 166 | **GGAAGCCCGATCCTCTTTTGAGGAACAAGTTTTCTTGTAAGACTTCAA** |
| 167 | **ATATCGCGTTTCCTCTTTTGAGGAACAAGTTTTCTTGTTTAATTCGAG** |
| 168 | **CTTCAAAGCGTCCTCTTTTGAGGAACAAGTTTTCTTGTAACCAGACCG** |
| 169 | **GAAGCAAACTTCCTCTTTTGAGGAACAAGTTTTCTTGTCCAACAGGTC** |
| 170 | **AGGATTAGAGTCCTCTTTTGAGGAACAAGTTTTCTTGTAGTACCTTTA** |
| 171 | **ATTGCTCCTTTCCTCTTTTGAGGAACAAGTTTTCTTGTTTGATAAGAG** |
| 172 | **GTCATTTTTGTCCTCTTTTGAGGAACAAGTTTTCTTGTCGGATGGCTT** |
| 173 | **AGAGCTTAATTCCTCTTTTGAGGAACAAGTTTTCTTGTTGCTGAATAT** |
| 174 | **AATGCTGTAGTCCTCTTTTGAGGAACAAGTTTTCTTGTCTCAACATGT** |
| 175 | **TTTAAATATGTCCTCTTTTGAGGAACAAGTTTTCTTGTCAACTAAAGT** |
| 176 | **ACGGTGTCTGTCCTCTTTTGAGGAACAAGTTTTCTTGTGAAGTTTCAT** |
| 177 | **TCCATATAACTCCTCTTTTGAGGAACAAGTTTTCTTGTAGTTGATTCC** |
| 178 | CAATTCTGCGAACGAGTAGATTTAGT |
| 179 | TTGACCATTAGATACATTTCGCAAATGGTCAATAACCT |
| 180 | GTTTAGCTATATTTTCATTTGGGGCGCGAGCTGAAAAG |
| 181 | GTGGCATCAATTCTACTAATAGTAGTAGCATTAACATC |
| 182 | CAATAAATCATACAGGCAAGGCAAAGAATTAGCAAAAT |
| 183 | TAAGCAATAAAGCCTCAGAGCATAAAGCTAAATCGGTT |
| 184 | GTACCAAAAACATTATGACCCTGTAATACTTTTGCGGG |
| 185 | AGAAGCCTTTATTTCAACGCAAGGATAAAAATTTTTAG |
| 186 | AACCCTCATATATTTTAAATGCAATGCCTGAGTAATGT |
| 187 | GTAGGTAAAGATTCAAAAGGGTGAGAAAGGCCGGAGAC |
| 188 | AGTCAAATCACCATCAATATGATATTCAACCGTTCTAG |
| 189 | CTGATAAATTAATGCCGGAGAGGGTAGCTATTTTTGAG |
| 190 | AGATCTACAAAGGCTATCAGGTCATTGCCTGAGAGTCT |
| 191 | GGAGCAAACAAGAGAATCGATGAACGGTAATCGTAAAA |
| 192 | CTAGCATGTCAATCATATGTACCCCGGTTGATAATCAG |
| 193 | AAAAGCCCCAAAAACAGGAAGATTGTATAAGCAAATAT |
| 194 | TTAAATTGTAAACGTTAATATTTTGTTAAAATTCGCAT |
| 195 | TAAATTTTTGTTAAATCAGCTCATTTTTTAACCAATAG |
| 196 | GAACGCCATCAAAAATAATTCGCGTCTGGCCTTCCTGT |
| 197 | AGCCAGCTTTCATCAACATTAAATGTGAGCGAGTAACA |
| 198 | ACCCGTCGGATTCTCCGTGGGAACAAACGGCGGATTGA |
| 199 | CCGTAATGGGATAGGTCACGTTGGTGTAGATGGGCGCA |
| 200 | TCGTAACCGTGCATCTGCCAGTTTGAGGGGACGACGAC |
| 201 | AGTATCGGCCTCAGGAAGATCGCACTCCAGCCAGCTTT |
| 202 | CCGGCACCGCTTCTGGTGCCGGAAACCAGGCAAAGCGC |
| 203 | CATTCGCCATTCAGGCTGCGCAACTGTTGGGAAGGGCG |
| 204 | ATCGGTGCGGGCCTCTTCGCTATTACGCCAGCTGGCGA |

| | |
|---|---|
| 205 | AAGGGGGATGTGCTGCAAGGCGATTAAGTTGGGTAACG |
| 206 | CCAGGGTTTTCCCAGTCACGACGTTGTAAAACGACGGC |
| 207 | CAGTGCCAAGCTTGCATGCCTGCAGGTCGACTCTAGAGGATCTTTT |

### Table S5.

Sequence of 1) biotinylated RNA aptamer.

| Oligo number | Sequence (5′→ 3′) |
|---|---|
| 1 | 3′-modification CGGAAUCAGUGAAUGCUUAUACAUCCGU-biotin |

Sequence of DNA oligonucleotides used for assembly of dsDNA nanostructure used for capture of biotinylated aptamer 'S'.

Oligonucleotide / **Dumbbell** / **Overhang**

| Oligo number | Sequence (5′→ 3′) |
|---|---|
| 1 | TTTTCGTAATCATGGTCATAGCTGTTTCCTGTGTGAAATTGTTATC |
| 2 | CGCTCACAATTCCACACAACATACGAGCCGGAAGCATA |
| 3 | AAGTGTAAAGCCTGGGGTGCCTAATGAGTGAGCTAACT |
| 4 | CACATTAATTGCGTTGCGCTCACTGCCCGCTTTCCAGT |
| 5 | CGGGAAACCTGTCGTGCCAGCTGCATTAATGAATCGGC |
| 6 | CAACGCGCGGGGAGAGGCGGTTTGCGTATTGGGCGCCA |
| 7 | GGGTGGTTTTTCTTTTCACCAGTGAGACGGGCAACAGC |
| 8 | TGATTGCCCTTCACCGCCTGGCCCTGAGAGAGTTGCAG |
| 9 | CAAGCGGTCCACGCTGGTTTGCCCCAGCAGGCGAAAAT |
| 10 | CCTGTTTGATGGTGGTTCCGAAATCGGCAAAATCCCTT |
| 11 | ATAAATCAAAAGAATAGCCCGAGATAGGGTTGAGTGTT |
| 12 | GTTCCAGTTTGGAACAAGAGTCCACTATTAAAGAACGT |
| 13 | GGACTCCAACGTCAAAGGGCGAAAAACCGTCTATCAGG |
| 14 | GCGATGGCCCACTACGTGAACCATCACCCAAATCAAGT |
| 15 | TTTTTGGGGTCGAGGTGCCGTAAAGCACTAAATCGGAA |
| 16 | CCCTAAAGGGAGCCCCCGATTTAGAGCTTGACGGGGAA |
| 17 | AGCCGGCGAACGTGGCGAGAAAGGAAGGGAAGAAAGCG |
| 18 | AAAGGAGCGGGCGCTAGGGCGCTGGCAAGTGTAGCGGT |
| 19 | CACGCTGCGCGTAACCACCACACCCGCCGCGCTTAATG |
| 20 | CGCCGCTACAGGGCGCGTACTATGGTTGCTTTGACGAG |
| 21 | CACGTATAACGTGCTTTCCTCGTTAGAATCAGAGCGGG |
| 22 | AGCTAAACAGGAGGCCGATTAAAGGGATTTTAGACAGG |
| 23 | AACGGTACGCCAGAATCCTGAGAAGTGTTTTTATAATC |
| 24 | AGTGAGGCCACCGAGTAAAAGAGTCTGTCCATCACGCA |
| 25 | AATTAACCGTTGTAGCAATACTTCTTTGATTAGTAATA |
| 26 | **ACATCACTTGTCCTCTTTTGAGGAACAAGTTTTCTTGTCCTGAGTAGA** |

| | |
|---|---|
| 27 | **AGAACTCAAATCCTCTTTTGAGGAACAAGTTTTCTTGTCTATCGGCCT** |
| 28 | **TGCTGGTAATTCCTCTTTTGAGGAACAAGTTTTCTTGTATCCAGAACA** |
| 29 | **ATATTACCGCTCCTCTTTTGAGGAACAAGTTTTCTTGTCAGCCATTGC** |
| 30 | **AACAGGAAAATCCTCTTTTGAGGAACAAGTTTTCTTGTACGCTCATGG** |
| 31 | **AAATACCTACTCCTCTTTTGAGGAACAAGTTTTCTTGTATTTTGACGC** |
| 32 | **TCAATCGTCTTCCTCTTTTGAGGAACAAGTTTTCTTGTGAAATGGATT** |
| 33 | **ATTTACATTGTCCTCTTTTGAGGAACAAGTTTTCTTGTGCAGATTCAC** |
| 34 | **CAGTCACACGTCCTCTTTTGAGGAACAAGTTTTCTTGTACCAGTAATA** |
| 35 | **AAAGGGACATTCCTCTTTTGAGGAACAAGTTTTCTTGTTCTGGCCAAC** |
| 36 | **AGAGATAGAATCCTCTTTTGAGGAACAAGTTTTCTTGTCCCTTCTGAC** |
| 37 | CTGAAAGCGTAAGAATACGTGGCACAGACAATATTTTTGAATGGCT |
| 38 | ATTAGTCTTTAATGCGCGAACTGATAGCCCTAAAACAT |
| 39 | CGCCATTAAAAATACCGAACGAACCACCAGCAGAAGAT |
| 40 | AAAACAGAGGTGAGGCGGTCAGTATTAACACCGCCTGC |
| 41 | AACAGTGCCACGCTGAGAGCCAGCAGCAAATGAAAAAT |
| 42 | CTAAAGCATCACCTTGCTGAACCTCAAATATCAAACCC |
| 43 | TCAATCAATATCTGGTCAGTTGGCAAATCAACAGTTGA |
| 44 | AAGGAATTGAGGAAGGTTATCTAAAATATCTTTAGGAG |
| 45 | AATATAATCCTGATTGTTTGGATTATACTTCTGAATAATGGAAGGG |
| 46 | **CACTAACAACTCCTCTTTTGAGGAACAAGTTTTCTTGTTAATAGATTA** |
| 47 | **GAGCCGTCAATCCTCTTTTGAGGAACAAGTTTTCTTGTTAGATAATAC** |
| 48 | **ATTTGAGGATTCCTCTTTTGAGGAACAAGTTTTCTTGTTTAGAAGTAT** |
| 49 | **TAGACTTTACTCCTCTTTTGAGGAACAAGTTTTCTTGTAAACAATTCG** |
| 50 | **ACAACTCGTATCCTCTTTTGAGGAACAAGTTTTCTTGTTTAAATCCTT** |
| 51 | **TGCCCGAACGTCCTCTTTTGAGGAACAAGTTTTCTTGTTTATTAATTT** |
| 52 | **TAAAAGTTTGTCCTCTTTTGAGGAACAAGTTTTCTTGTAGTAACATTA** |
| 53 | **TCATTTTGCGTCCTCTTTTGAGGAACAAGTTTTCTTGTGAACAAAGAA** |
| 54 | **ACCACCAGAATCCTCTTTTGAGGAACAAGTTTTCTTGTGGAGCGGAAT** |
| 55 | **TATCATCATATCCTCTTTTGAGGAACAAGTTTTCTTGTTTCCTGATTA** |
| 56 | **TCAGATGATGTCCTCTTTTGAGGAACAAGTTTTCTTGTGCAATTCATC** |
| 57 | TTAGAACCTACCATATCAAAATTATTTGCACGTAAAAC |
| 58 | AGAAATAAAGAAATTGCGTAGATTTTCAGGTTTAACGT |
| 59 | CAGATGAATATACAGTAACAGTACCTTTTACATCGGGA |
| 60 | GAAACAATAACGGATTCGCCTGATTGCTTTGAATACCA |
| 61 | AGTTACAAAATCGCGCAGAGGCGAATTATTCATTTCAA |
| 62 | TTACCTGAGCAAAAGAAGATGATGAAACAAACATCAAG |
| 63 | AAAACAAAATTAATTACATTTAACAATTTCATTTGAAT |
| 64 | TACCTTTTTAATGGAAACAGTACATAAATCAATATAT |
| 65 | GTGAGTGAATAACCTTGCTTCTGTAAATCGTCGCTATT |
| 66 | AATTAATTTTCCCTTAGAATCCTTGAAAACATAGCGAT |
| 67 | AGCTTAGATTAAGACGCTGAGAAGAGTCAATAGTGAAT |
| 68 | TTATCAAAATCATAGGTCTGAGAGACTACCTTTTTAAC |
| 69 | CTCCGGCTTAGGTTGGGTTATATAACTATATGTAAATG |

| | |
|---|---|
| 70 | CTGATGCAAATCCAATCGCAAGACAAAGAACGCGAGAA |
| 71 | AACTTTTTCAAATATATTTTAGTTAATTTCATCTTCTG |
| 72 | ACCTAAATTTAATGGTTTGAAATACCGACCGTGTGATA |
| 73 | AATAAGGCGTTAAATAAGAATAAACACCGGAATCATAA |
| 74 | TTACTAGAAAAAGCCTGTTTAGTATCATATGCGTTATA |
| 75 | CAAATTCTTACCAGTATAAAGCCAACGCTCAACAGTAG |
| 76 | GGCTTAATTGAGAATCGCCATATTTAACAACGCCAACA |
| 77 | TGTAATTTAGGCAGAGGCATTTTCGAGCCAGTAATAAG |
| 78 | AGAATATAAAGTACCGACAAAAGGTAAAGTAATTCTGT |
| 79 | CCAGACGACGACAATAAACAACATGTTCAGCTAATGCA |
| 80 | GAACGCGCCTGTTTATCAACAATAGATAAGTCCTGAAC |
| 81 | AAGAAAAATAATATCCCATCCTAATTTACGAGCATGTA |
| 82 | GAAACCAATCAATAATCGGCTGTCTTTCCTTATCATTC |
| 83 | CAAGAACGGGTATTAAACCAAGTACCGCACTCATCGAG |
| 84 | AACAAGCAAGCCGTTTTTATTTTCATCGTAGGAATCAT |
| 85 | TACCGCGCCCAATAGCAAGCAAATCAGATATAGAAGGC |
| 86 | TTATCCGGTATTCTAAGAACGCGAGGCGTTTTAGCGAA |
| 87 | CCTCCCGACTTGCGGGAGGTTTTGAAGCCTTAAATCAA |
| 88 | GATTAGTTGCTATTTTGCACCCAGCTACAATTTTATCC |
| 89 | TGAATCTTACCAACGCTAACGAGCGTCTTTCCAGAGCC |
| 90 | TAATTTGCCAGTTACAAAATAAACAGCCATATTATTTA |
| 91 | TCCCAATCCAAATAAGAAACGATTTTTTGTTTAACGTC |
| 92 | AAAAATGAAAATAGCAGCCTTTACAGAGAGAATAACAT |
| 93 | AAAAACAGGGAAGCGCATTAGACGGGAGAATTAACTGA |
| 94 | ACACCCTGAACAAAGTCAGAGGGTAATTGAGCGCTAAT |
| 95 | GAATTGAGTTAAGCCCAA |
| 96 | **ACGGATGTATAAGCATTCACTGATTCCGTTTTTATCAGAGAGATAACCCACAA** |
| 97 | TAATAAGAGCAAGAAACAATGAAATAGCAATAGCTATC |
| 98 | TTACCGAAGCCCTTTTTAAGAAAAGTAAGCAGATAGCC |
| 99 | GAACAAAGTTACCAGAAGGAAACCGAGGAAACGCAATA |
| 100 | ATAACGGAATACCCAAAAGAACTGGCATGATTAAGACT |
| 101 | CCTTATTACGCAGTATGTTAGCAAACGTAGAAAATACA |
| 102 | TACATAAAGGTGGCAACATATAAAAGAAACGCAAAGAC |
| 103 | ACCACGGAATAAGTTTATTTTGTCACAATCAATAGAAA |
| 104 | ATTCATATGGTTTACCAGCGCCAAAGACAAAAGGGCGA |
| 105 | CATTCAACCGATTGAGGGAGGGAAGGTAAATATTGACG |
| 106 | GAAATTATTCATTAAAGGTGAATTATCACCGTCACCGA |
| 107 | CTTGAGCCATTTGGGAATTAGAGCCAGCAAAATCACCA |
| 108 | GTAGCACCATTACCATTAGCAAGGCCGGAAACGTCACC |
| 109 | AATGAAACCATCGATAGCAGCACCGTAATCAGTAGCGA |
| 110 | CAGAATCAAGTTTGCCTTTAGCGTCAGACTGTAGCGCG |
| 111 | TTTTCATCGGCATTTTCGGTCATAGCCCCCTTATTAGC |
| 112 | GTTTGCCATCTTTTCATAATCAAAATCACCGGAACCAG |

| | |
|---|---|
| 113 | AGCCACCACCGGAACCGCCTCCCTCAGAGCCGCCACCC |
| 114 | TCAGAACCGCCACCCTCAGAGCCACCACCCTCAGAGCC |
| 115 | GCCACCAGAACCACCACCAGAGCCGCCGCCAGCATTGA |
| 116 | CAGGAGGTTGAGGCAGGTCAGACGATTGGCCTTGATAT |
| 117 | TCACAAACAAATAAATCCTCATTAAAGCCAGAATGGAA |
| 118 | AGCGCAGTCTCTGAATTTACCGTTCCAGTAAGCGTCAT |
| 119 | ACATGGCTTTTGATGATACAGGAGTGTACTGGTAATAA |
| 120 | GTTTTAACGGGGTCAGTGCCTTGAGTAACAGTGCCCGT |
| 121 | ATAAACAGTTAATGCCCCCTGCCTATTTCGGAACCTAT |
| 122 | TATTCTGAAACATGAAAGTATTAAGAGGCTGAGACTCC |
| 123 | TCAAGAGAAGGATTAGGATTAGCGGGGTTTTGCTCAGT |
| 124 | ACCAGGCGGATAAGTGCCGTCGAGAGGGTTGATATAAG |
| 125 | TATAGCCCGGAATAGGTGTATCACCGTACTCAGGAGGT |
| 126 | TTAGTACCGCCACCCTCAGAACCGCCACCCTCAGAACC |
| 127 | GCCACCCTCAGAGCCACCACCCTCATTTTCAGGGATAG |
| 128 | CAAGCCCAATAGGAACCCATGTACCGTAACACTGAGTT |
| 129 | TCGTCACCAGTACAAACTACAACGCCTGTAGCATTCCA |
| 130 | CAGACAGCCCTCATAGTTAGCGTAACGATCTAAAGTTT |
| 131 | TGTCGTCTTTCCAGACGTTAGTAAATGAATTTTCTGTA |
| 132 | TGGGATTTTGCTAAACAACTTTCAACAGTTTCAGCGGA |
| 133 | GTGAGAATAGAAAGGAACAACTAAAGGAATTGCGAATA |
| 134 | ATAATTTTTTCACGTTGAAAATCTCCAAAAAAAAGGCT |
| 135 | CCAAAAGGAGCCTTTAATTGTATCGGTTTATCAGCTTG |
| 136 | CTTTCGAGGTGAATTTCTTAAACAGCTTGATACCGATA |
| 137 | GTTGCGCCGACAATGACAACAACCATCGCCCACGCATA |
| 138 | ACCGATATATTCGGTCGCTGAGGCTTGCAGGGAGTTAA |
| 139 | AGGCCGCTTTTGCGGGATCGTCACCCTCAGCAGCGAAA |
| 140 | GACAGCATCGGAACGAGGGTAGCAACGGCTACAGAGGC |
| 141 | TTTGAGGACTAAAGACTTTTTCATGAGGAAGTTTCCAT |
| 142 | TAAACGGGTAAAATACGTAATGCCACTACGAAGGCACC |
| 143 | AACCTAAAACGAAAGAGGCAAAAGAATACACTAAAACA |
| 144 | CTCATCTTTGACCCCCAGCGATTATACCAAGCGCGAAA |
| 145 | CAAAGTACAACGGAGATTTGTATCATCGCCTGATAAAT |
| 146 | TGTGTCGAAATCCGCGACCTGCTCCATGTTACTTAGCC |
| 147 | GGAACGAGGCGCAGACGGTCAATCATAAGGGAACCGAA |
| 148 | CTGACCAACTTTGAAAGAGGACAGATGAACGGTGTACA |
| 149 | GACCAGGCGCATAGGCTGGCTGACCTTCATCAAGAGTA |
| 150 | ATCTTGACAAGAACCGGATATTCATTACCCAAATCAAC |
| 151 | GTAACAAAGCTGCTCATTCAGTGAATAAGGCTTGCCCT |
| 152 | GACGAGAAACACCAGAACGAGTAGTAAATTGGGCTTGA |
| 153 | GATGGTTTAATTTCAACTTTAATCATTGTGAATTACCT |
| 154 | TATGCGATTTTAAGAACTGGCTCATTATACCAGTCAGG |
| 155 | ACGTTGGGAAGAAAAATCTACGTTAATAAAACGAACTA |

| | |
|---|---|
| 156 | ACGGAACAACATTATTACAGGTAGAAAGATTCATCAGT |
| 157 | TGAGATTTAGGAATACCACATTCAACTAATGCAGATAC |
| 158 | ATAACGCCAAAAGGAATTACGAGGCATAGTAAGAGCAA |
| 159 | CACTATCATAACCCTCGTTTACCAGACGACGATAAAAA |
| 160 | CCAAAATAGCGAGAGGCTTTTGCAAAAGAAGTTTTGCC |
| 161 | AGAGGGGGTAATAGTAAAATGTTTAGACTGGATAGCGT |
| 162 | CCAATACTGCGGAATCGTCATAAATATTCATTGAATCC |
| 163 | CCCTCAAATGCTTTAAACAGTTCAGAAAACGAGAATGA |
| 164 | CCATAAATCAAAAATCAGGTCTTTACCCTGACTATTAT |
| 165 | AGTCAGAAGCAAAGCGGATTGCATCAAAAAGATTAAGA |
| 166 | **GGAAGCCCGATCCTCTTTTGAGGAACAAGTTTTCTTGTAAGACTTCAA** |
| 167 | **ATATCGCGTTTCCTCTTTTGAGGAACAAGTTTTCTTGTTTAATTCGAG** |
| 168 | **CTTCAAAGCGTCCTCTTTTGAGGAACAAGTTTTCTTGTAACCAGACCG** |
| 169 | **GAAGCAAACTTCCTCTTTTGAGGAACAAGTTTTCTTGTCCAACAGGTC** |
| 170 | **AGGATTAGAGTCCTCTTTTGAGGAACAAGTTTTCTTGTAGTACCTTTA** |
| 171 | **ATTGCTCCTTTCCTCTTTTGAGGAACAAGTTTTCTTGTTTGATAAGAG** |
| 172 | **GTCATTTTTGTCCTCTTTTGAGGAACAAGTTTTCTTGTCGGATGGCTT** |
| 173 | **AGAGCTTAATTCCTCTTTTGAGGAACAAGTTTTCTTGTTGCTGAATAT** |
| 174 | **AATGCTGTAGTCCTCTTTTGAGGAACAAGTTTTCTTGTCTCAACATGT** |
| 175 | **TTTAAATATGTCCTCTTTTGAGGAACAAGTTTTCTTGTCAACTAAAGT** |
| 176 | **ACGGTGTCTGTCCTCTTTTGAGGAACAAGTTTTCTTGTGAAGTTTCAT** |
| 177 | **TCCATATAACTCCTCTTTTGAGGAACAAGTTTTCTTGTAGTTGATTCC** |
| 178 | CAATTCTGCGAACGAGTAGATTTAGT |
| 179 | TTGACCATTAGATACATTTCGCAAATGGTCAATAACCT |
| 180 | GTTTAGCTATATTTTCATTTGGGGCGCGAGCTGAAAAG |
| 181 | GTGGCATCAATTCTACTAATAGTAGTAGCATTAACATC |
| 182 | CAATAAATCATACAGGCAAGGCAAAGAATTAGCAAAAT |
| 183 | TAAGCAATAAAGCCTCAGAGCATAAAGCTAAATCGGTT |
| 184 | GTACCAAAAACATTATGACCCTGTAATACTTTTGCGGG |
| 185 | AGAAGCCTTTATTTCAACGCAAGGATAAAAATTTTTAG |
| 186 | AACCCTCATATATTTTAAATGCAATGCCTGAGTAATGT |
| 187 | GTAGGTAAAGATTCAAAAGGGTGAGAAAGGCCGGAGAC |
| 188 | AGTCAAATCACCATCAATATGATATTCAACCGTTCTAG |
| 189 | CTGATAAATTAATGCCGGAGAGGGTAGCTATTTTTGAG |
| 190 | AGATCTACAAAGGCTATCAGGTCATTGCCTGAGAGTCT |
| 191 | GGAGCAAACAAGAGAATCGATGAACGGTAATCGTAAAA |
| 192 | CTAGCATGTCAATCATATGTACCCCGGTTGATAATCAG |
| 193 | AAAAGCCCCAAAAACAGGAAGATTGTATAAGCAAATAT |
| 194 | TTAAATTGTAAACGTTAATATTTTGTTAAAATTCGCAT |
| 195 | TAAATTTTTGTTAAATCAGCTCATTTTTTAACCAATAG |
| 196 | GAACGCCATCAAAAATAATTCGCGTCTGGCCTTCCTGT |
| 197 | AGCCAGCTTTCATCAACATTAAATGTGAGCGAGTAACA |
| 198 | ACCCGTCGGATTCTCCGTGGGAACAAACGGCGGATTGA |

| | |
|---|---|
| 199 | CCGTAATGGGATAGGTCACGTTGGTGTAGATGGGCGCA |
| 200 | TCGTAACCGTGCATCTGCCAGTTTGAGGGGACGACGAC |
| 201 | AGTATCGGCCTCAGGAAGATCGCACTCCAGCCAGCTTT |
| 202 | CCGGCACCGCTTCTGGTGCCGGAAACCAGGCAAAGCGC |
| 203 | CATTCGCCATTCAGGCTGCGCAACTGTTGGGAAGGGCG |
| 204 | ATCGGTGCGGGCCTCTTCGCTATTACGCCAGCTGGCGA |
| 205 | AAGGGGGATGTGCTGCAAGGCGATTAAGTTGGGTAACG |
| 206 | CCAGGGTTTTCCCAGTCACGACGTTGTAAAACGACGGC |
| 207 | CAGTGCCAAGCTTGCATGCCTGCAGGTCGACTCTAGAGGATCTTTT |

### Table S6.

Sequence of oligonucleotides attached to 1) 0, 2) 5, 3) 10, 4) 20 and 40 kDA PEG used in nanostructure comparing PEG molecular weight (Figure 4a-d).

| Oligo number | Sequence (5′→ 3′) |
|---|---|
| 1 | GTCTGGGCTGGATGTTGTCG |
| 2 | CTGATCCTAGTGACGAAACC-5′5 kDa PEG |
| 3 | ACCACTAATGAGTGATATCC-5′10 kDa PEG |
| 4 | TGAGGATGTGTAGGTTGAGG-5′20 kDa PEG |
| 5 | CGGAAUCAGUGAAUGCUUAUACAUCCGU -5′40 kDa PEG (RNA) |

Sequence of DNA oligonucleotide mixture binding 20 and 40 kDA PEG (Figure 4e-f).

| Oligo number | Sequence (5′→ 3′) |
|---|---|
| 1 | TGAGGATGTGTAGGTTGAGG -5′20 kDa PEG |
| 2 | TGAGGATGTGTAGGTTGAGG -5′40 kDa PEG |

### Table S7.

Sequence of DNA oligonucleotides used for assembly of dsDNA nanostructure used for capture of PEGylated oligonucleotide conjugates.

Oligonucleotide / **Dumbbell** / **Overhangs**

| Oligo number | Sequence (5′→ 3′) |
|---|---|
| 1 | TTTTCGTAATCATGGTCATAGCTGTTTCCTGTGTGAAATTGTTATC |
| 2 | CGCTCACAATTCCACACAACATACGAGCCGGAAGCATA |
| 3 | AAGTGTAAAGCCTGGGGTGCCTAATGAGTGAGCTAACT |
| 4 | CACATTAATTGCGTTGCGCTCACTGCCCGCTTTCCAGT |
| 5 | CGGGAAACCTGTCGTGCCAGCTGCATTAATGAATCGGC |
| 6 | CAACGCGCGGGGAGAGGCGGTTTGCGTATTGGGCGCCA |
| 7 | GGGTGGTTTTTCTTTTCACCAGTGAGACGGGCAACAGC |
| 8 | TGATTGCCCTTCACCGCCTGGCCCTGAGAGAGTTGCAG |
| 9 | CAAGCGGTCCACGCTGGTTTGCCCCAGCAGGCGAAAAT |
| 10 | CCTGTTTGATGGTGGTTCCGAAATCGGCAAAATCCCTT |
| 11 | ATAAATCAAAAGAATAGCCCGAGATAGGGTTGAGTGTT |
| 12 | GTTCCAGTTTGGAACAAGAGTCCACTATTAAAGAACGT |
| 13 | GGACTCCAACGTCAAAGGGCGAAAAACCGTCTATCAGG |
| 14 | GCGATGGCCCACTACGTGAACCATCACCCAAATCAAGT |
| 15 | TTTTTGGGGTCGAGGTGCCGTAAAGCACTAAATCGGAA |
| 16 | CCCTAAAGGGAGCCCCCGATTTAGAGCTTGACGGGGAA |
| 17 | AGCCGGCGAACGTGGCGAGAAAGGAAGGGAAGAAAGCG |
| 18 | AAAGGAGCGGGCGCTAGGGCGCTGGCAAGTGTAGCGGT |
| 19 | CACGCTGCGCGTAACCACCACACCCGCCGCGCTTAATG |
| 20 | CGCCGCTACAGGGCGCGTACTATGGTTGCTTTGACGAG |
| 21 | CACGTATAACGTGCTTTCCTCGTTAGAATCAGAGCGGG |
| 22 | AGCTAAACAGGAGGCCGATTAAAGGGATTTTAGACAGG |
| 23 | AACGGTACGCCAGAATCCTGAGAAGTGTTTTTATAATC |
| 24 | AGTGAGGCCACCGAGTAAAAGAGTCTGTCCATCACGCA |
| 25 | AATTAACCGTTGTAGCAATACTTCTTTGATTAGTAATA |
| 26 | **ACATCACTTGTCCTCTTTTGAGGAACAAGTTTTCTTGTCCTGAGTAGA** |
| 27 | **AGAACTCAAATCCTCTTTTGAGGAACAAGTTTTCTTGTCTATCGGCCT** |
| 28 | **TGCTGGTAATTCCTCTTTTGAGGAACAAGTTTTCTTGTATCCAGAACA** |
| 29 | **ATATTACCGCTCCTCTTTTGAGGAACAAGTTTTCTTGTCAGCCATTGC** |
| 30 | **AACAGGAAAATCCTCTTTTGAGGAACAAGTTTTCTTGTACGCTCATGG** |
| 31 | **AAATACCTACTCCTCTTTTGAGGAACAAGTTTTCTTGTATTTTGACGC** |
| 32 | **TCAATCGTCTTCCTCTTTTGAGGAACAAGTTTTCTTGTGAAATGGATT** |

| | |
|---|---|
| 33 | **ATTTACATTGTCCTCTTTTGAGGAACAAGTTTTCTTGTGCAGATTCAC** |
| 34 | **CAGTCACACGTCCTCTTTTGAGGAACAAGTTTTCTTGTACCAGTAATA** |
| 35 | **AAAGGGACATTCCTCTTTTGAGGAACAAGTTTTCTTGTTCTGGCCAAC** |
| 36 | **AGAGATAGAATCCTCTTTTGAGGAACAAGTTTTCTTGTCCCTTCTGAC** |
| 37 | CTGAAAGCGTAAGAATACGTGGCACAGACAATATTTTTGAATGGCT |
| 38 | ATTAGTCTTTAATGCGCGAACTGATAGCCCTAAAACAT |
| 39 | CGCCATTAAAAATACCGAACGAACCACCAGCAGAAGAT |
| 40 | AAAACAGAGGTGAGGCGGTCAGTATTAACACCGCCTGC |
| 41 | AACAGTGCCACGCTGAGAGCCAGCAGCAAATGAAAAAT |
| 42 | CTAAAGCATCACCTTGCTGAACCTCAAATATCAAACCC |
| 43 | TCAATCAATATCTGGTCAGTTGGCAAATCAACAGTTGA |
| 44 | AAGGAATTGAGGAAGGTTATCTAAAATATCTTTAGGAG |
| 45 | CACTAACAACTAATAGATTAGAGCCGTCAATAGATAAT |
| 46 | ACATTTGAGGATTTAGAAGTATTAGACTTTACAAACAA |
| 47 | TTCGACAACTCGTATTAAATCCTTTGCCCGAACGTTAT |
| 48 | TAATTTTAAAAGTTTGAGTAACATTATCATTTTGCGGA |
| 49 | **cgacaacatccagcccagactttttACAAAGAAACCACCAGAAGG** |
| 50 | AGCGGAATTATCATCATA |
| 51 | TTCCTGATTATCAGATGATGGCAATTCATCAATATAAT |
| 52 | CCTGATTGTTTGGATTATACTTCTGAATAATGGAAGGG |
| 53 | TTAGAACCTACCATATCAAAATTATTTGCACGTAAAAC |
| 54 | AGAAATAAAGAAATTGCGTAGATTTTCAGGTTTAACGT |
| 55 | CAGATGAATATACAGTAACAGTACCTTTTACATCGGGA |
| 56 | GAAACAATAACGGATTCGCCTGATTGCTTTGAATACCA |
| 57 | AGTTACAAAATCGCGCAGAGGCGAATTATTCATTTCAA |
| 58 | TTACCTGAGCAAAAGAAGATGATGAAACAAACATCAAG |
| 59 | AAAACAAAATTAATTACATTTAACAATTTCATTTGAAT |
| 60 | TACCTTTTTTAATGGAAACAGTACATAAATCAATATAT |
| 61 | GTGAGTGAATAACCTTGCTTCTGTAAATCGTCGCTATT |
| 62 | **ggtttcgtcactaggatcagtttttAGCTTAGATTAAGACGCTGA** |
| 63 | GAAGAGTCAATAGTGAAT |
| 64 | TTATCAAAATCATAGGTCTGAGAGACTACCTTTTTAAC |
| 65 | CTCCGGCTTAGGTTGGGTTATATAACTATATGTAAATG |
| 66 | CTGATGCAAATCCAATCGCAAGACAAAGAACGCGAGAA |
| 67 | AACTTTTTCAAATATATTTTAGTTAATTTCATCTTCTG |
| 68 | ACCTAAATTTAATGGTTTGAAATACCGACCGTGTGATA |
| 69 | AATAAGGCGTTAAATAAGAATAAACACCGGAATCATAA |
| 70 | TTACTAGAAAAAGCCTGTTTAGTATCATATGCGTTATA |
| 71 | CAAATTCTTACCAGTATAAAGCCAACGCTCAACAGTAG |
| 72 | GGCTTAATTGAGAATCGCCATATTTAACAACGCCAACA |
| 73 | TGTAATTTAGGCAGAGGCATTTTCGAGCCAGTAATAAG |
| 74 | AGAATATAAAGTACCGACAAAAGGTAAAGTAATTCTGT |
| 75 | **GGATATCACTCATTAGTGGTtttttGAACGCGCCTGTTTATCAAC** |

| | |
|---|---|
| 76 | AATAGATAAGTCCTGAAC |
| 77 | AAGAAAAATAATATCCCATCCTAATTTACGAGCATGTA |
| 78 | GAAACCAATCAATAATCGGCTGTCTTTCCTTATCATTC |
| 79 | CAAGAACGGGTATTAAACCAAGTACCGCACTCATCGAG |
| 80 | AACAAGCAAGCCGTTTTTATTTTCATCGTAGGAATCAT |
| 81 | TACCGCGCCCAATAGCAAGCAAATCAGATATAGAAGGC |
| 82 | TTATCCGGTATTCTAAGAACGCGAGGCGTTTTAGCGAA |
| 83 | CCTCCCGACTTGCGGGAGGTTTTGAAGCCTTAAATCAA |
| 84 | GATTAGTTGCTATTTTGCACCCAGCTACAATTTTATCC |
| 85 | TGAATCTTACCAACGCTAACGAGCGTCTTTCCAGAGCC |
| 86 | TAATTTGCCAGTTACAAAATAAACAGCCATATTATTTA |
| 87 | TCCCAATCCAAATAAGAAACGATTTTTTGTTTAACGTC |
| 88 | **cctcaacctacacatcctcatttttAAAAACAGGGAAGCGCATTA** |
| 89 | GACGGGAGAATTAACTGA |
| 90 | ACACCCTGAACAAAGTCAGAGGGTAATTGAGCGCTAAT |
| 91 | ATCAGAGAGATAACCCACAAGAATTGAGTTAAGCCCAA |
| 92 | TAATAAGAGCAAGAAACAATGAAATAGCAATAGCTATC |
| 93 | TTACCGAAGCCCTTTTTAAGAAAAGTAAGCAGATAGCC |
| 94 | GAACAAAGTTACCAGAAGGAAACCGAGGAAACGCAATA |
| 95 | ATAACGGAATACCCAAAAGAACTGGCATGATTAAGACT |
| 96 | CCTTATTACGCAGTATGTTAGCAAACGTAGAAAATACA |
| 97 | TACATAAAGGTGGCAACATATAAAAGAAACGCAAAGAC |
| 98 | ACCACGGAATAAGTTTATTTTGTCACAATCAATAGAAA |
| 99 | ATTCATATGGTTTACCAGCGCCAAAGACAAAAGGGCGA |
| 100 | CATTCAACCGATTGAGGGAGGGAAGGTAAATATTGACG |
| 101 | **CTTGAGCCATTTGGGAATTAttttttACGGATGTATAAGCATTCACTGATTCCG** |
| 102 | GAGCCAGCAAAATCACCA |
| 103 | GTAGCACCATTACCATTAGCAAGGCCGGAAACGTCACC |
| 104 | AATGAAACCATCGATAGCAGCACCGTAATCAGTAGCGA |
| 105 | CAGAATCAAGTTTGCCTTTAGCGTCAGACTGTAGCGCG |
| 106 | TTTTCATCGGCATTTTCGGTCATAGCCCCCTTATTAGC |
| 107 | GTTTGCCATCTTTTCATAATCAAAATCACCGGAACCAG |
| 108 | AGCCACCACCGGAACCGCCTCCCTCAGAGCCGCCACCC |
| 109 | TCAGAACCGCCACCCTCAGAGCCACCACCCTCAGAGCC |
| 110 | GCCACCAGAACCACCACCAGAGCCGCCGCCAGCATTGA |
| 111 | CAGGAGGTTGAGGCAGGTCAGACGATTGGCCTTGATAT |
| 112 | TCACAAACAAATAAATCCTCATTAAAGCCAGAATGGAA |
| 113 | AGCGCAGTCTCTGAATTTACCGTTCCAGTAAGCGTCAT |
| 114 | ACATGGCTTTTGATGATACAGGAGTGTACTGGTAATAA |
| 115 | **GTTTTAACGGTCCTCTTTTGAGGAACAAGTTTTCTTGTGGTCAGTGCC** |
| 116 | **TTGAGTAACATCCTCTTTTGAGGAACAAGTTTTCTTGTGTGCCCGTAT** |
| 117 | **AAACAGTTAATCCTCTTTTGAGGAACAAGTTTTCTTGTTGCCCCCTGC** |
| 118 | **CTATTTCGGATCCTCTTTTGAGGAACAAGTTTTCTTGTACCTATTATT** |

| | |
|---|---|
| 119 | **CTGAAACATGTCCTCTTTTGAGGAACAAGTTTTCTTGTAAAGTATTAA** |
| 120 | **GAGGCTGAGATCCTCTTTTGAGGAACAAGTTTTCTTGTCTCCTCAAGA** |
| 121 | **GAAGGATTAGTCCTCTTTTGAGGAACAAGTTTTCTTGTGATTAGCGGG** |
| 122 | **GTTTTGCTCATCCTCTTTTGAGGAACAAGTTTTCTTGTGTACCAGGCG** |
| 123 | **GATAAGTGCCTCCTCTTTTGAGGAACAAGTTTTCTTGTGTCGAGAGGG** |
| 124 | **TTGATATAAGTCCTCTTTTGAGGAACAAGTTTTCTTGTTATAGCCCGG** |
| 125 | **AATAGGTGTATCCTCTTTTGAGGAACAAGTTTTCTTGTTCACCGTACT** |
| 126 | CAGGAGGTTTAGTACCGCCACCCTCAGAACCGCCACCCTCAGAACC |
| 127 | GCCACCCTCAGAGCCACCACCCTCATTTTCAGGGATAG |
| 128 | CAAGCCCAATAGGAACCCATGTACCGTAACACTGAGTT |
| 129 | TCGTCACCAGTACAAACTACAACGCCTGTAGCATTCCA |
| 130 | CAGACAGCCCTCATAGTTAGCGTAACGATCTAAAGTTT |
| 131 | TGTCGTCTTTCCAGACGTTAGTAAATGAATTTTCTGTA |
| 132 | TGGGATTTTGCTAAACAACTTTCAACAGTTTCAGCGGA |
| 133 | GTGAGAATAGAAAGGAACAACTAAAGGAATTGCGAATA |
| 134 | ATAATTTTTTCACGTTGAAAATCTCCAAAAAAAAGGCT |
| 135 | CCAAAAGGAGCCTTTAATTGTATCGGTTTATCAGCTTG |
| 136 | CTTTCGAGGTGAATTTCTTAAACAGCTTGATACCGATA |
| 137 | GTTGCGCCGACAATGACAACAACCATCGCCCACGCATA |
| 138 | ACCGATATATTCGGTCGCTGAGGCTTGCAGGGAGTTAA |
| 139 | AGGCCGCTTTTGCGGGATCGTCACCCTCAGCAGCGAAA |
| 140 | GACAGCATCGGAACGAGGGTAGCAACGGCTACAGAGGC |
| 141 | TTTGAGGACTAAAGACTTTTTCATGAGGAAGTTTCCAT |
| 142 | TAAACGGGTAAAATACGTAATGCCACTACGAAGGCACC |
| 143 | AACCTAAAACGAAAGAGGCAAAAGAATACACTAAAACA |
| 144 | CTCATCTTTGACCCCCAGCGATTATACCAAGCGCGAAA |
| 145 | CAAAGTACAACGGAGATTTGTATCATCGCCTGATAAAT |
| 146 | TGTGTCGAAATCCGCGACCTGCTCCATGTTACTTAGCC |
| 147 | GGAACGAGGCGCAGACGGTCAATCATAAGGGAACCGAA |
| 148 | CTGACCAACTTTGAAAGAGGACAGATGAACGGTGTACA |
| 149 | GACCAGGCGCATAGGCTGGCTGACCTTCATCAAGAGTA |
| 150 | ATCTTGACAAGAACCGGATATTCATTACCCAAATCAAC |
| 151 | GTAACAAAGCTGCTCATTCAGTGAATAAGGCTTGCCCT |
| 152 | GACGAGAAACACCAGAACGAGTAGTAAATTGGGCTTGA |
| 153 | GATGGTTTAATTTCAACTTTAATCATTGTGAATTACCT |
| 154 | TATGCGATTTTAAGAACTGGCTCATTATACCAGTCAGG |
| 155 | ACGTTGGGAAGAAAAATCTACGTTAATAAAACGAACTA |
| 156 | ACGGAACAACATTATTACAGGTAGAAAGATTCATCAGT |
| 157 | TGAGATTTAGGAATACCACATTCAACTAATGCAGATAC |
| 158 | ATAACGCCAAAAGGAATTACGAGGCATAGTAAGAGCAA |
| 159 | CACTATCATAACCCTCGTTTACCAGACGACGATAAAAA |
| 160 | CCAAAATAGCGAGAGGCTTTTGCAAAAGAAGTTTTGCC |
| 161 | AGAGGGGGTAATAGTAAAATGTTTAGACTGGATAGCGT |

| | |
|---|---|
| 162 | CCAATACTGCGGAATCGTCATAAATATTCATTGAATCC |
| 163 | CCCTCAAATGCTTTAAACAGTTCAGAAAACGAGAATGA |
| 164 | CCATAAATCAAAAATCAGGTCTTTACCCTGACTATTAT |
| 165 | AGTCAGAAGCAAAGCGGATTGCATCAAAAAGATTAAGA |
| 166 | GGAAGCCCGAAAGACTTCAAATATCGCGTTTTAATTCG |
| 167 | AGCTTCAAAGCGAACCAGACCGGAAGCAAACTCCAACA |
| 168 | GGTCAGGATTAGAGAGTACCTTTAATTGCTCCTTTTGA |
| 169 | TAAGAGGTCATTTTTGCGGATGGCTTAGAGCTTAATTG |
| 170 | CTGAATATAATGCTGTAGCTCAACATGTTTTAAATATG |
| 171 | CAACTAAAGTACGGTGTCTGGAAGTTTCATTCCATATA |
| 172 | ACAGTTGATTCCCAATTCTGCGAACGAGTAGATTTAGT |
| 173 | TTGACCATTAGATACATTTCGCAAATGGTCAATAACCT |
| 174 | GTTTAGCTATATTTTCATTTGGGGCGCGAGCTGAAAAG |
| 175 | GTGGCATCAATTCTACTAATAGTAGTAGCATTAACATC |
| 176 | CAATAAATCATACAGGCAAGGCAAAGAATTAGCAAAAT |
| 177 | TAAGCAATAAAGCCTCAGAGCATAAAGCTAAATCGGTT |
| 178 | GTACCAAAAACATTATGACCCTGTAATACTTTTGCGGG |
| 179 | AGAAGCCTTTATTTCAACGCAAGGATAAAAATTTTTAG |
| 180 | AACCCTCATATATTTTAAATGCAATGCCTGAGTAATGT |
| 181 | GTAGGTAAAGATTCAAAAGGGTGAGAAAGGCCGGAGAC |
| 182 | AGTCAAATCACCATCAATATGATATTCAACCGTTCTAG |
| 183 | CTGATAAATTAATGCCGGAGAGGGTAGCTATTTTTGAG |
| 184 | AGATCTACAAAGGCTATCAGGTCATTGCCTGAGAGTCT |
| 185 | GGAGCAAACAAGAGAATCGATGAACGGTAATCGTAAAA |
| 186 | CTAGCATGTCAATCATATGTACCCCGGTTGATAATCAG |
| 187 | AAAAGCCCCAAAAACAGGAAGATTGTATAAGCAAATAT |
| 188 | TTAAATTGTAAACGTTAATATTTTGTTAAAATTCGCAT |
| 189 | TAAATTTTTGTTAAATCAGCTCATTTTTTAACCAATAG |
| 190 | GAACGCCATCAAAAATAATTCGCGTCTGGCCTTCCTGT |
| 191 | AGCCAGCTTTCATCAACATTAAATGTGAGCGAGTAACA |
| 192 | ACCCGTCGGATTCTCCGTGGGAACAAACGGCGGATTGA |
| 193 | CCGTAATGGGATAGGTCACGTTGGTGTAGATGGGCGCA |
| 194 | TCGTAACCGTGCATCTGCCAGTTTGAGGGGACGACGAC |
| 195 | AGTATCGGCCTCAGGAAGATCGCACTCCAGCCAGCTTT |
| 196 | CCGGCACCGCTTCTGGTGCCGGAAACCAGGCAAAGCGC |
| 197 | CATTCGCCATTCAGGCTGCGCAACTGTTGGGAAGGGCG |
| 198 | ATCGGTGCGGGCCTCTTCGCTATTACGCCAGCTGGCGA |
| 199 | AAGGGGGATGTGCTGCAAGGCGATTAAGTTGGGTAACG |
| 200 | CCAGGGTTTTCCCAGTCACGACGTTGTAAAACGACGGC |
| 201 | CAGTGCCAAGCTTGCATGCCTGCAGGTCGACTCTAGAGGATCTTTT |

### Table S8.

Statistical analysis for current spike observables in **Figures 4c-4d**, including descriptive statistics, Shapiro-Wlik test, Levene's test, OMNIBUS Test, and post-hoc Games-Howell pairwise comparisons.

***Figure 4c***

**Descriptive Statistics**

| Group | n | Mean | Std | Median | Min | Max |
|---|---|---|---|---|---|---|
| 0 kDa | 194 | 0.1168 | 0.0511 | 0.1061 | 0.0217 | 0.3642 |
| 5 kDa | 194 | 0.1756 | 0.0613 | 0.1832 | 0.0276 | 0.3371 |
| 10 kDa | 194 | 0.2191 | 0.0776 | 0.2306 | 0.0360 | 0.4712 |
| 20 kDa | 194 | 0.2409 | 0.1009 | 0.2653 | 0.0335 | 0.5372 |
| 40 kDa | 194 | 0.4317 | 0.1422 | 0.4702 | 0.0367 | 0.6521 |

**Normality: Shapiro-Wilk test ($H_0$: data are normal)**

| Group | W | p-value | Normality |
|---|---|---|---|
| 0 kDa | 0.9256 | 0.0000 | No |
| 5 kDa | 0.9870 | 0.0712 | Yes |
| 10 kDa | 0.9757 | 0.0019 | No |
| 20 kDa | 0.9133 | 0.0000 | No |
| 40 kDa | 0.8637 | 0.0000 | No |

**→ All groups normal: False**

**Homogeneity of variance: Levene's test ($H_0$: variances equal)**

- Levene statistic: 24.8335
- p-value: 0.0000 ***
- Equal variances: False

**OMNIBUS Test**

- **Welch's ANOVA**
  $F(4, 471.1300) = 255.1412$, $p = 0.000000$ ***
  Significant: True
- **Standard ANOVA (for reference)**
  $F = 319.8750$, $p = 0.000000$ ***
- **Kruskal-Wallis (non-parametric)**
  $H = 453.3789$, $p = 0.000000$ ***
  Significant: True

**Post-hoc — Games-Howell pairwise comparisons**

| Pair | D | SE | q | df | p-value |
|---|---|---|---|---|---|
| 0-5 | -0.0588 | 0.0057 | 14.5155 | 374.0970 | 0.0000 |
| 0-10 | -0.1023 | 0.0067 | 21.6725 | 333.9870 | 0.0000 |
| 0-20 | -0.1241 | 0.0081 | 21.6080 | 285.9970 | 0.0000 |
| 0-40 | -0.3149 | 0.0108 | 41.0480 | 242.1309 | 0.0000 |
| 5-10 | -0.0435 | 0.0071 | 8.6613 | 366.1833 | 0.0000 |
| 5-20 | -0.0653 | 0.0085 | 10.8975 | 318.1897 | 0.0000 |
| 5-40 | -0.2561 | 0.0111 | 32.5812 | 262.2579 | 0.0000 |
| 10-20 | -0.0218 | 0.0091 | 3.3777 | 362.1608 | 0.0072 |
| 10-40 | -0.2126 | 0.0116 | 25.8497 | 298.6876 | 0.0000 |
| 20-40 | -0.1908 | 0.0125 | 21.5496 | 348.1219 | 0.0000 |

**Summary**

- Groups compared: 5
  Equal variances: False (Levene p=0.0000)
  All groups normal: False

- Suitable test: Welch's ANOVA
  Suitable post-hoc: Games-Howell

- Welch's ANOVA: F=255.1412., p=0.000000 ***
  Kruskal-Wallis: H=255.1412, p=0.000000 ***

***Figure 4d***

**Descriptive Statistics**

| Group | n | Mean | Std | Median | Min | Max |
|---|---|---|---|---|---|---|
| 0 kDa | 194 | 4.81 | 1.68 | 4.56 | 2.09 | 11.41 |
| 5 kDa | 194 | 6.50 | 2.01 | 6.59 | 2.04 | 11.07 |
| 10 kDa | 194 | 7.31 | 2.45 | 7.05 | 2.37 | 20.04 |
| 20 kDa | 194 | 9.24 | 3.56 | 9.23 | 2.14 | 23.32 |
| 40 kDa | 194 | 13.20 | 3.88 | 13.80 | 2.63 | 26.39 |

**Normality: Shapiro-Wilk test ($H_0$: data are normal)**

| Group | W | p-value | Normality |
|---|---|---|---|
| 0 kDa | 0.9503 | 0.0000 | No |
| 5 kDa | 0.9906 | 0.2404 | Yes |
| 10 kDa | 0.9505 | 0.0000 | No |
| 20 kDa | 0.9809 | 0.0095 | No |
| 40 kDa | 0.9599 | 0.0000 | No |

**→ All groups normal: False**

**Homogeneity of variance: Levene's test ($H_0$: variances equal)**

- Levene statistic: 26.9254
- p-value: 0.0000 ***
- Equal variances: False

**OMNIBUS Test**

- **Welch's ANOVA**
  F(4, 472.24) = 224.2523, p = 0.000000 ***
  Significant: True
- **Standard ANOVA (for reference)**
  F = 245.5839, p = 0.000000 ***
- **Kruskal-Wallis (non-parametric)**
  H = 451.9016, p = 0.000000 ***
  Significant: True

**Post-hoc — Games-Howell pairwise comparisons**

| Pair | D | SE | q | df | p-value |
|---|---|---|---|---|---|
| 0-5 | -1.7022 | 0.1885 | 12.7706 | 374.4255 | 0.0000 |
| 0-10 | -2.5075 | 0.2138 | 16.5839 | 341.8132 | 0.0000 |
| 0-20 | -4.4430 | 0.2834 | 22.1744 | 275.0080 | 0.0000 |
| 0-40 | -8.3949 | 0.3041 | 39.0427 | 263.1407 | 0.0000 |
| 5-10 | -0.8053 | 0.2280 | 4.9959 | 371.7029 | 0.0000 |
| 5-20 | -2.7409 | 0.2942 | 13.1762 | 304.5259 | 0.0000 |
| 5-40 | -6.6927 | 0.3142 | 30.1253 | 289.6456 | 0.0000 |
| 10-20 | -1.9355 | 0.3110 | 8.8008 | 342.2837 | 0.0000 |
| 10-40 | -5.8874 | 0.3300 | 25.2297 | 325.9556 | 0.0000 |
| 20-40 | -3.9519 | 0.3788 | 14.7546 | 383.2421 | 0.0000 |

**Summary**

- Groups compared: 5
  Equal variances: False  (Levene p=0.0000)
  All groups normal: False

- Suitable test: Welch's ANOVA
  Suitable post-hoc: Games-Howell

- Welch's ANOVA: F=224.2523, p=0.000000 ***
  Kruskal-Wallis: H=451.9016,  p=0.000000 ***

**Table S9.**

The table includes the IV curves of the nanopores used to characterize the DNA nanostructures prior to the measurement. The ionic current and RMS noise at 600 mV is presented for each pore presented in the Main text.

| **Pore number** | **Sample** | **Current, RMS noise at 600 mV** | **Current / voltage curve (IV curve)** |
|---|---|---|---|
| 1 | DNA nanostructures 'P' and 'R' (Figure 1 and Figure 2a-c) | 12.6 nA<br>6.1 pA | 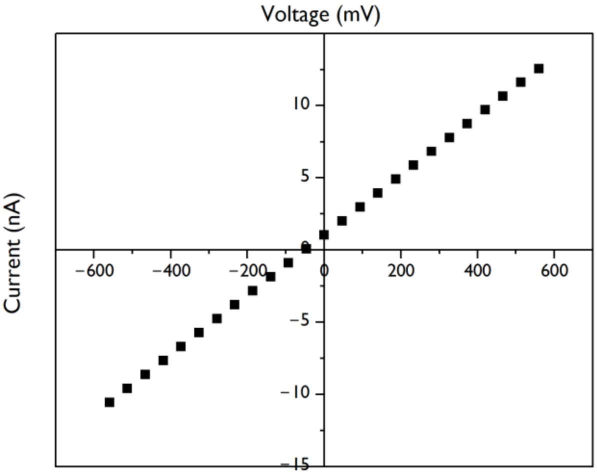  |
| 2 | DNA nanostructure 'P' in horse serum (Figure 2d-f) | 8.4 nA<br>6.3 pA | 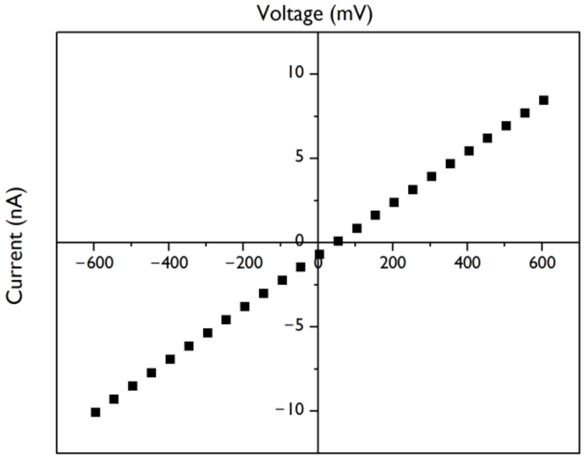  |

| | | | |
|---|---|---|---|
| 3 | DNA nanostructure ‘S’ (Figure 3a-d) | 7.7 nA<br>6.5 pA | 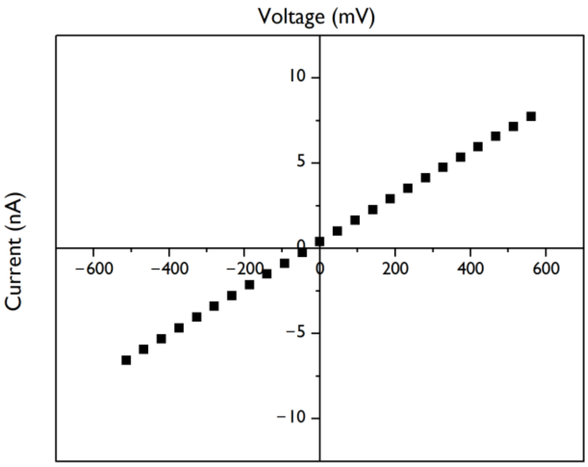 |
| 4 | DNA nanostructure ‘S’ and ‘P’ 5min in horse serum (Figure 3e) | 9.1 nA<br>5.8 pA | 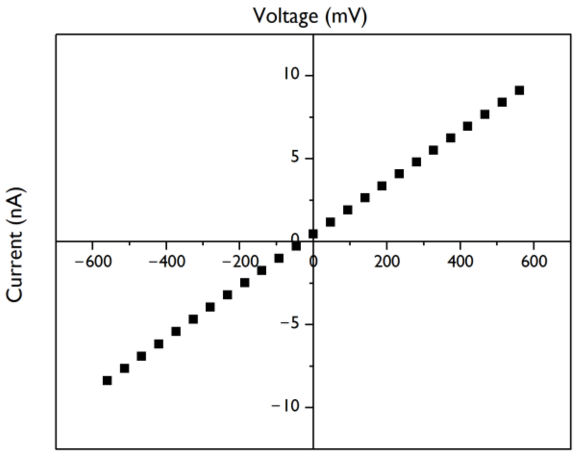 |
| 5 | DNA nanostructure ‘S’ and ‘P’ 24 h in horse serum (Figure 3f) | 12.9 nA<br>6.4 pA | 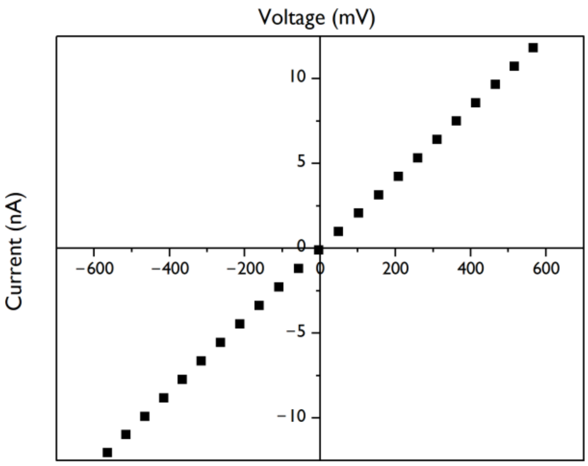 |

| | | | |
|---|---|---|---|
| 6 | DNA nanostructure for 0,5, 10, 20 kDa PEG and competing capture nanostructure (Figure 4) | 8.1 nA<br>6.4 pA | 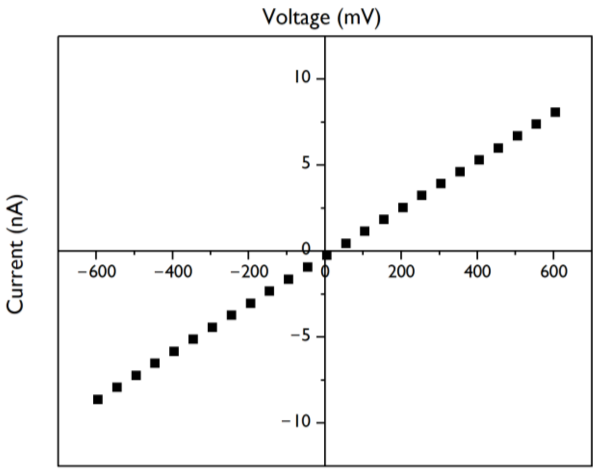 |
